# Biomass segregation between biofilm and flocs improves the control of nitrite-oxidizing bacteria in mainstream partial nitritation and anammox processes

**DOI:** 10.1101/480780

**Authors:** Michele Laureni, David G. Weissbrodt, Kris Villez, Orlane Robin, Nadieh de Jonge, Alex Rosenthal, George Wells, Jeppe Lund Nielsen, Eberhard Morgenroth, Adriano Joss

## Abstract

The control of nitrite-oxidizing bacteria (NOB) challenges the implementation of partial nitritation and anammox (PN/A) processes under mainstream conditions. The aim of the present study was to understand how operating conditions impact microbial competition and the control of NOB in hybrid PN/A systems, where biofilm and flocs coexist. A hybrid PN/A moving-bed biofilm reactor (MBBR; also referred to as integrated fixed film activated sludge or IFAS) was operated at 15 °C on aerobically pre-treated municipal wastewater (23 mg_NH4-N_·L^−1^). Ammonium-oxidizing bacteria (AOB) and NOB were enriched primarily in the flocs, and anammox bacteria (AMX) in the biofilm. After decreasing the dissolved oxygen concentration (DO) from 1.2 to 0.17 mg_O2_·L^−1^ - with all other operating conditions unchanged - washout of NOB from the flocs was observed. The activity of the minor NOB fraction remaining in the biofilm was suppressed at low DO. As a result, low effluent NO_3_^−^ concentrations (0.5 mg_N_·L^−1^) were consistently achieved at aerobic nitrogen removal rates (80 mg_N_·L^−1^·d^−1^) comparable to those of conventional treatment plants. A simple dynamic mathematical model, assuming perfect biomass segregation with AOB and NOB in the flocs and AMX in the biofilm, was able to qualitatively reproduce the selective washout of NOB from the flocs in response to the decrease in DO-setpoint. Similarly, numerical simulations indicated that flocs removal is an effective operational strategy to achieve the selective washout of NOB. The direct competition for NO_2_^−^ between NOB and AMX - the latter retained in the biofilm and acting as a “NO_2_-sink” - was identified by the model as key mechanism leading to a difference in the actual growth rates of AOB and NOB (*i.e*., μ_NOB_ < μ_AOB_ in flocs) and allowing for the selective NOB washout. Experimental results and model predictions demonstrate the increased operational flexibility, in terms of variables that can be easily controlled by operators, offered by hybrid systems as compared to solely biofilm systems for the control of NOB in mainstream PN/A applications.

**Highlights:**

- Hybrid PN/A systems provide increased operational flexibility for NOB control
- AOB and NOB enrich primarily in the flocs, and AMX in the biofilm (“NO_2_-sink”)
- AMX use NO_2_^−^ allowing to differentiate AOB and NOB growth rates
- A decrease in DO or an increase in floc removal leads to selective NOB washout from flocs
- The activity of the minor NOB fraction in the biofilm is suppressed at limiting DO

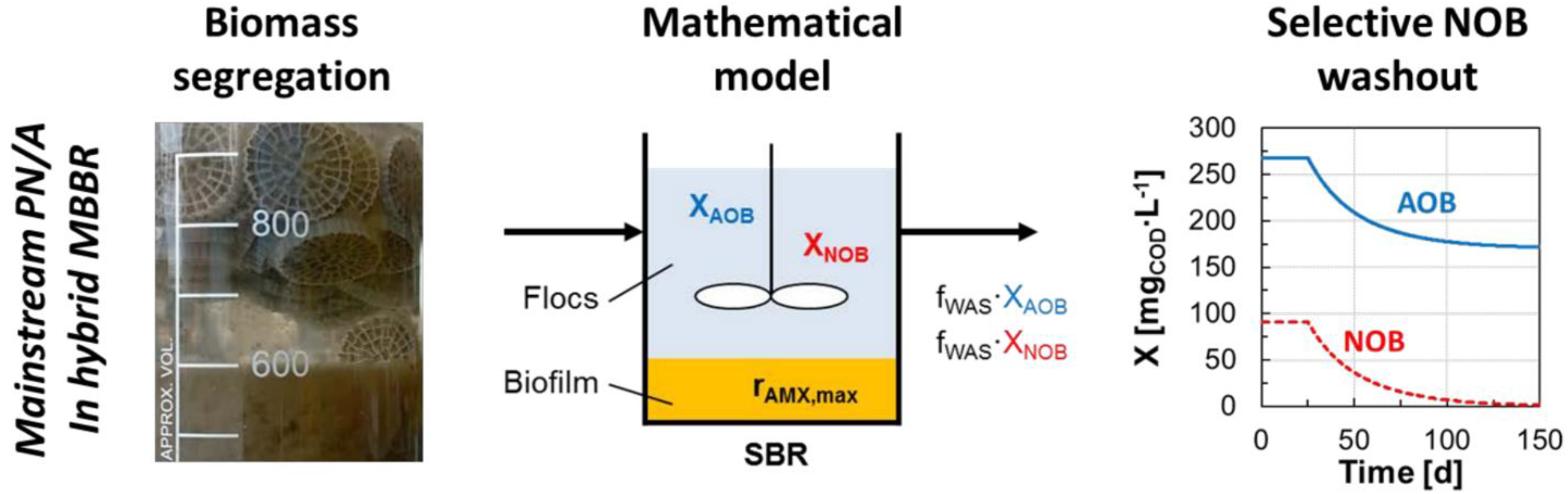

## 1 Introduction

Partial nitritation and anammox (PN/A) is a resource-efficient alternative process for the removal of nitrogen from municipal wastewater (MWW) and holds promise to bring wastewater treatment plants (WWTP) close to neutral or even positive energy balances (Siegrist *et al*., 2008, van Loosdrecht and Brdjanovic 2014). PN/A technologies are implemented for the treatment of warm and concentrated streams such as digester supernatant (“sidestream PN/A”; Lackner *et al*., (2014)). Research targeting the direct application of PN/A to more dilute MWW, or “mainstream PN/A”, is progressing at a fast pace (De Clippeleir *et al*., 2013, Gilbert *et al*., 2015a, Laureni *et al*., 2016, Lotti *et al*., 2015). The challenges associated with mainstream PN/A relate to the highly variable, dilute and cold characteristics of MWW. Moreover, mainstream PN/A must guarantee volumetric N-removal rates comparable to conventional WWTP (*i.e*., 100 mg_N_·L^−1^d^−1^; Metcalf & Eddy *et al*., (2013)) and reliably discharge effluent to stringent water quality standards (e.g., below 2 mg_NH4-N_·L^−1^ in Switzerland; WPO (1998)).

Successful PN/A relies on the concerted activity of aerobic (AOB) and anaerobic ammonium-oxidizing (AMX) bacteria (Speth *et al*., 2016). Optimized microbial community engineering strategies are required to favour the growth of AOB and retain the slower-growing AMX, while out-competing the undesired nitrite-oxidizing bacteria (NOB). Several operational strategies implemented in sidestream applications are not feasible under mainstream conditions. At mesophilic temperatures (> 20°C), AOB display higher maximum growth rates than NOB, which allows selective NOB washout at a sufficiently low solids retention time. Conversely, at mainstream temperatures between 10-20°C (in temperate regions), the differences in growth rates are minimal (Hellinga *et al*., 1998). In addition, nitrogen concentrations in the main line are too low for NOB to be inhibited by free ammonia (NH3) or free nitrous acid (HNO_2_) (Anthonisen *et al*., 1976, Jubany *et al*., 2009). As a result, NOB control and washout cannot be based on maximum growth rates alone, as is efficiently achieved in sidestream suspended biomass systems (Hellinga *et al*., 1998, Joss *et al*., 2011).

The use of biofilms, either grown on carrier material or in the form of granular bio-aggregates, has proven effective to achieve stable and resilient PN/A under mainstream conditions at laboratory scale (Gilbert *et al*., 2015a, Laureni *et al*., 2016, Lotti *et al*., 2015). Biofilms allow for the long solids retention times (SRT) needed to retain AMX, while substrate gradients promote the suppression of NOB activity (Brockmann and Morgenroth 2010, Gilbert *et al*., 2015a, Laureni *et al*., 2016, Lotti *et al*., 2014, Pérez *et al*., 2014). NOB control in biofilm systems is mainly driven by the competition for oxygen with AOB, with the latter usually featuring higher substrate affinities (Brockmann and Morgenroth 2010, Corbala-Robles *et al*., 2016, Pérez *et al*., 2014). PN/A operation under oxygen-limited NH4+ oxidation can favour nitritation while limiting the aerobic growth of NOB (Brockmann and Morgenroth 2010, Isanta *et al*., 2015, Pérez *et al*., 2014). However, operation under oxygen limitation inherently limits the AOB activity as well, and thus the overall process rate (Laureni *et al*., 2015, Perez *et al*., 2014). Moreover, despite the generally accepted higher affinity of AOB for oxygen (Rittmann and McCarty 2001), NOB are known to adapt to low dissolved oxygen concentrations (DO) (Liu and Wang 2013), and several studies have recently reported higher oxygen affinities for NOB than AOB (Malovanyy *et al*., 2015, Regmi *et al*., 2014, Sliekers *et al*., 2005). Lastly, although their activity can be suppressed, NOB can persist in the biofilm and become active when favourable conditions are re-established, making their long-term suppression in solely biofilm systems challenging (Fux *et al*., 2004, Gilbert *et al*., 2015a, Isanta *et al*., 2015, Laureni *et al*., 2016, Lotti *et al*., 2014).

Hybrid systems, where biofilms and flocs coexist (also referred to as integrated fixed film activated sludge or IFAS), are currently receiving increased attention for their potential advantages for PN/A applications. Experimental evidence (Laureni *et al*., 2016, Leix *et al*., 2016, Malovanyy *et al*., 2015, Park *et al*., 2014, Shi *et al*., 2016, Veuillet *et al*., 2014, Vlaeminck *et al*., 2010, Wells *et al*., 2017, Winkler *et al*., 2011) and numerical results (Hubaux *et al*., 2015, Volcke *et al*., 2012) indicate that the faster-growing aerobic guilds tend to enrich in the floc fraction, with direct access to dissolved substrates. In turn, AMX have been shown to enrich in the biofilm, where anoxic conditions are achieved. As a result, differential control of the retention times of the bacterial guilds associated with the two biomass fractions is in principle possible (Wett *et al*., 2015). Moreover, as flocs are less diffusion-limited than biofilms, significantly higher aerobic volumetric conversion rates can be achieved even at low DO (Veuillet *et al*., 2014). Nonetheless, published data on hybrid systems operated for PN/A remain limited and seemingly contradictory. Hybrid systems at high flocs concentrations above 1 g_TSS_·L^−1^ have been applied at full scale to treat digester supernatant at mesophilic temperatures with negligible NOB activity (Veuillet *et al*., 2014). Conversely, increased NOB activity has been reported in hybrid systems with a fraction of flocs as small as < 10% of total solids (Hubaux *et al*., 2015, Laureni *et al*., 2016). The implications of biomass segregation and operational conditions for microbial competition in hybrid systems are as yet largely unknown.

The aim of this work was to understand the dominant mechanisms controlling the interaction between biofilm and flocs, the influence of operating conditions, and their implications for NOB control in hybrid PN/A systems. The effect of the DO on NOB was assessed experimentally in an IFAS system operated on real MWW at 15°C. In parallel, a simplified dynamic mathematical model of the hybrid system was developed to provide a mechanistic interpretation of the experimental results, and to understand how the composition of the flocs and the NOB concentration respond to changes in DO, flocs removal, and AMX activity in the biofilm. The sensitivity of the simulation outcome to model parameters was assessed. Relevant scenarios for engineering practice are also discussed.

## 2 Materials and methods

### 2.1 Long-term reactor operation at different DO

A 12 L hybrid MBBR was operated as a sequencing batch reactor (SBR) for PN/A on aerobically pre-treated MWW (see next section). The reactor was filled at a volumetric ratio of 33% with K5 biofilm carriers (AnoxKaldnes^™^, Sweden; protected surface of 800 m^2^·m^−3^). The biomass was previously acclimatised to the influent for over one year (Laureni *et al*., 2016). The reactor was run for 565 days at 15.5 ± 1.0°C. Each SBR cycle consisted of six steps: feeding (5 L of pre-treated MWW, 5 min), anoxic mixing (10 min; 200 rpm), aeration and mixing (variable duration terminated at a residual NH_4_^+^ concentration of 2 mg_NH4-N_·L^−1^), anoxic mixing (60 min), settling (60 min), and effluent discharge (terminated at 7 L fill level; 2 min). The DO was varied between micro-aerobic conditions (*Phases I, III, V:* 0.17 ± 0.04 mg_O2_·L^−1^; (Gilbert *et al*., 2015b)), and aerobic conditions (*Phases II, IV*: 1.2 ± 0.2 mg_O2_·L^−1^ and 1.6 ± 0.1 mg_O2_·L^−1^; (Regmi *et al*., 2014)) (Figure 2). The total cycle duration varied between 3.5 ± 0.5 and 5.3 ± 0.3 h for operation at high and low DO, respectively.

The reactor was equipped with an optical oxygen sensor (Oxymax COS61D), ion-selective electrodes for NH4+ and NO_3_^−^ concentrations, and pH and temperature sensors (ISEmax CAS40D), all from Endress+Hauser (Switzerland). The pH was not controlled and remained stable at 7.4 ± 0.2 throughout the experimental period. Operational data are presented in Figure S1.

### 2.2 Municipal wastewater (MWW)

The municipal wastewater was taken from the sewer of Dübendorf (Switzerland). After primary treatment (screen, sand removal and primary clarifier), MWW was pre-treated in an aerated 12 L SBR operated for high-rate organic carbon (as COD) removal at an SRT of 1 d. The pre-treated MWW featured the following characteristics: 54 ± 13 mg_CODsol_·L^−1^, 23 ± 6 mg_NH4-N_·L^−1^, and < 0.3 mg_N_·L^−1^ of NO_2_^−^ and NO_3_^−^. Prior to feeding to the PN/A reactor, the pre-treated MWW was stored in a temperature-controlled (< 20°C) external buffer tank of 50 L to equalize the hydraulic loads.

### 2.3 Control of total suspended solids (TSS) and calculation of their dynamic SRT

In addition to the settling step in the SBR cycle, from day 70 onwards the reactor effluent was filtered through a 10 L filter-bag (50-μm-mesh; 3M^™^ NB Series, Nylon Monofilament) placed in a 50 L barrel. The content of the net was centrifuged for 5 min at 2000 × g, and the solids were reintroduced into the reactor on a daily basis. The TSS in the reactor and all activities were measured one cycle after biomass reintroduction.

The dynamic total SRT was calculated considering only the observed sludge loss in the effluent and by sampling (modified from Takács *et al*., (2008)):

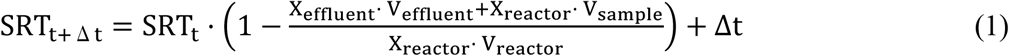

where X_effluent_ is the average TSS concentration in the sock-net effluent (g_TSS_·L^−1^), V_effluent_ is the total effluent volume discharged during the time interval, V_sample_ is the volume taken out for biomass sampling, X_reactor_ is the TSS concentration in the reactor (g_TSS_·L^−1^), V_reactor_ is the volume of the bulk liquid phase in the reactor (12 L), and Δt is the time interval between subsequent measurements (d). The aerobic SRT is calculated from the total SRT as follows:

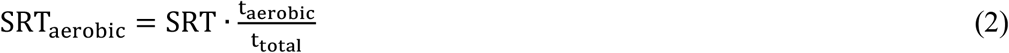

where t_aerobic_/t_total_ is the actual fraction of aerobic time over the total batch time (Figure S1). The development of TSS, SRT and SRT_aerobic_ over time is presented in Figure S2, together with the volumetric particle size distribution of the flocs measured on days 451 and 465 via laser light scattering (Mastersizer 2000, Malvern, UK).

### 2.4 Maximum activities of AOB, NOB and AMX, and their segregation between biofilm and flocs

The maximum anammox activity (r_AMX,max_) is defined as the volumetric rate of nitrogen removal (sum of NH_4_^+^ and NO_2_^−^) in the absence of DO and under non-limiting concentrations of NH_4_^+^ and NO_2_^−^. r_AMX,max_ was measured *in-situ* once or twice a week. The maximum activities of AOB and NOB (r_AOB,max_ and r_NOB,max_) are defined respectively as the volumetric rates of NH_4_^+^ oxidation and NO_3_^−^ production. r_AOB,max_ and r_NOB,max_ were measured via *ex-situ* batch tests (1 L) run under fully aerobic conditions (> 5 mgO_2_·L^−1^) and non-limiting concentrations of NH_4_^+^ and NO_2_^−^. The liquid fraction was sampled during mixing and a proportional number of random carriers were chosen manually. Mixing was provided with a magnetic stirrer (200 rpm) and the temperature was maintained at 15 ± 1°C. After manually removing all carriers, r_AOB,max_ and r_NOB,max_ of the flocs were measured. The rAMX,max value of the suspension was checked *ex-situ* five times throughout the experimental period and was confirmed to be negligible. NH_4_^+^ and NO_2_^−^ were supplied as NH_4_Cl and NaNO_2_ (20-30 mg_N_·L^−1^), and volumetric consumption rates were calculated by linear regression of laboratory measurements of 3-4 grab samples from the bulk liquid phase.

### 2.5 Activities of AOB, NOB, and AMX during regular operation (aerobic step)

The volumetric activities of the three main autotrophic guilds during regular operation (r_AOB,cycle_, r_NOB,cycle_ and r_AMX,cycle_ expressed as mg_NH4-N_·L^−1^·d^−1^, mg_NO3-N_·L^−1^·d^−1^, and mg_(NH4+NO2)-N_·L^−1^·d^−1^ respectively) were estimated according to Laureni *et al*., (2016). In short, during the aerated step of an SBR cycle, the consumption of NH_4_^+^, accumulation of NO_2_^−^ and production of NO_3_^−^ were followed by laboratory measurements of 3-4 grab samples from the bulk liquid phase. The activities were estimated based on the stoichiometric and kinetic matrix presented in Table 1, with parameters from Table 2. Heterotrophic denitrification during aeration was assumed to be negligible (Laureni *et al*., 2016).

**Table 1:**
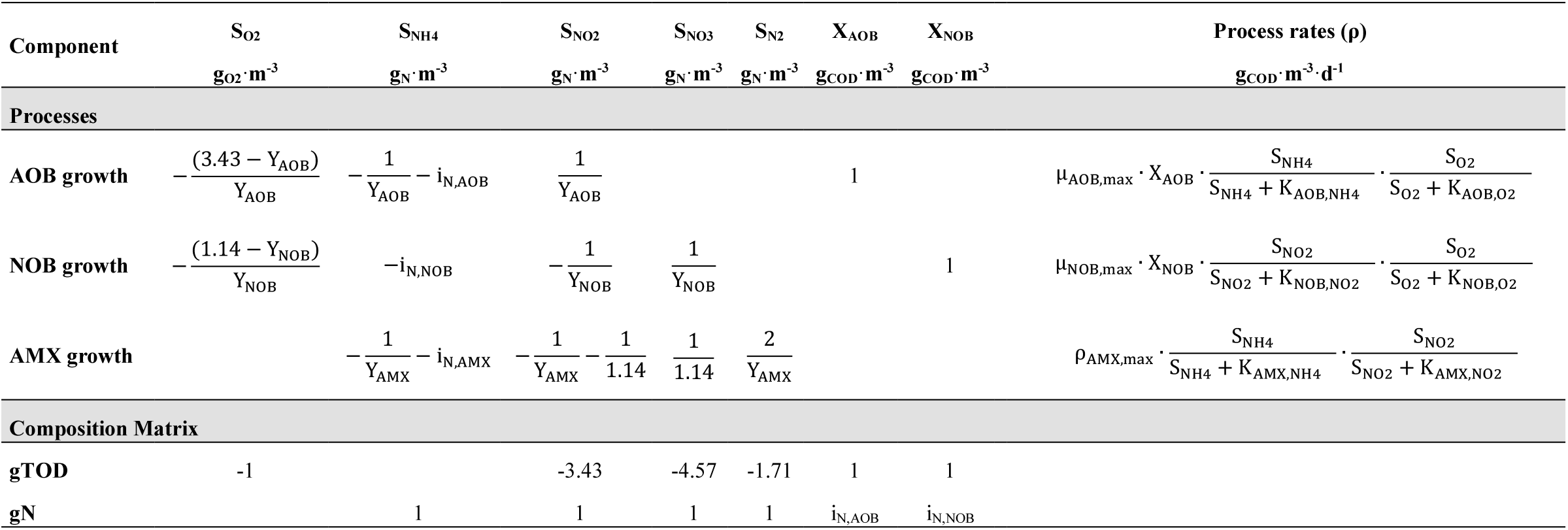
Stoichiometric and kinetic matrix describing the growth of aerobic ammonium-oxidizing bacteria (AOB) and aerobic nitrite-oxidizing bacteria (NOB), and anaerobic ammonium-oxidizing bacteria (anammox, AMX). The matrix was used to estimate the activity of the three guilds during regular SBR operation (r_i,cycle_), and for the dynamic model of the hybrid system (Figure 1). In the dynamic model, the maximum anammox process rate (ρ_AMX,max_= μ_AMX,max_·X_AMX_) was assumed constant during each simulation. To this end, the concentration of AMX (X_AMX_) was considered as a constant and not as a state variable, and is therefore omitted from the matrix.

**Table 2:** Kinetic and stoichiometric parameters.

| **Aerobic ammonium-oxidizing bacteria (AOB)** |  |  |  |  |
| --- | --- | --- | --- | --- |
| $\mu_{\text{AOB,max}}$ | $\text{d}^{-1}$ | Maximum specific growth rate | 0.30 | *This study\** |
| $Y_{\text{AOB}}$ | $\text{g}_{\text{COD}} \cdot \text{g}_{\text{N}}^{-1}$ | Growth yield | 0.18 | (Jubany *et al.*, 2009) |
| $K_{\text{NH}_4,\text{AOB}}$ | $\text{g}_{\text{NH}_4\text{-N}} \cdot \text{m}^{-3}$ | Ammonium half-saturation constant | 2.4 | (Wiesmann, 1994) |
| $K_{\text{O}_2,\text{AOB}}$ | $\text{g}_{\text{COD}} \cdot \text{m}^{-3}$ | Oxygen half-saturation constant | 0.6 | (Wiesmann, 1994) |
| $i_{\text{N,AOB}}$ | $\text{g}_{\text{N}} \cdot \text{g}_{\text{COD}}^{-1}$ | Nitrogen content in AOB | 0.083 | (Volcke *et al.*, 2010) |
| **Aerobic nitrite-oxidizing bacteria (NOB)** |  |  |  |  |
| $\mu_{\text{NOB,max}}$ | $\text{d}^{-1}$ | Maximum specific growth rate | 0.34 | *This study\** |
| $Y_{\text{NOB}}$ | $\text{g}_{\text{COD}} \cdot \text{g}_{\text{N}}^{-1}$ | Growth yield | 0.08 | (Jubany *et al.*, 2009) |
| $K_{\text{O}_2,\text{NOB}}$ | $\text{g}_{\text{COD}} \cdot \text{m}^{-3}$ | Oxygen half-saturation constant | 0.4 | (Blackburne *et al.*, 2007) |
| $K_{\text{NO}_2,\text{NOB}}$ | $\text{g}_{\text{NO}_2\text{-N}} \cdot \text{m}^{-3}$ | Nitrite half-saturation constant | 0.5 | (Wiesmann, 1994) |
| $i_{\text{N,NOB}}$ | $\text{g}_{\text{N}} \cdot \text{g}_{\text{COD}}^{-1}$ | Nitrogen content in NOB | 0.083 | (Volcke *et al.*, 2010) |
| **Anaerobic ammonium-oxidizing bacteria (AMX)** |  |  |  |  |
| $\rho_{\text{AMX,max}}$ | $\text{mg}_{\text{COD}} \cdot \text{L}^{-1} \cdot \text{d}^{-1}$ | Maximum AMX process rate | 0 - 24 | Assumed** |
| $Y_{\text{AMX}}$ | $\text{g}_{\text{COD}} \cdot \text{g}_{\text{N}}^{-1}$ | Growth yield | 0.17 | (Strous *et al.*, 1998) |
| $K_{\text{NH}_4,\text{AMX}}$ | $\text{g}_{\text{NH}_4\text{-N}} \cdot \text{m}^{-3}$ | Ammonium half saturation constant | 0.03 | (Volcke *et al.*, 2010) |
| $K_{\text{NO}_2,\text{AMX}}$ | $\text{g}_{\text{NO}_2\text{-N}} \cdot \text{m}^{-3}$ | Nitrite half saturation constant | 0.005 | (Volcke *et al.*, 2010) |
| $i_{\text{N,AMX}}$ | $\text{g}_{\text{N}} \cdot \text{g}_{\text{COD}}^{-1}$ | Nitrogen content in AMX | 0.058 | (Volcke *et al.*, 2010) |

\*Estimated from the maximum activity increase at 15°C during *Phase II* (Figure 2a).
\*\* Corresponding to $r_{\text{AMX,max}}$ in the range observed experimentally at 15°C, 0-300 $\text{mg}_{(\text{NH}_4+\text{NO}_2)\text{-N}} \cdot \text{L}^{-1} \cdot \text{d}^{-1}$

### 2.6 Nitrogen removal over the entire SBR cycle and during the aerobic step

Over the entire SBR cycle, the volumetric N-removal rate (mg_N_·L^−1^·d^−1^) was calculated by dividing the difference between the sum of the dissolved nitrogen species (NH_4_^+^, NO_2_^−^ and NO_3_^−^) in the influent and effluent by the hydraulic retention time (HRT, d). The relative removals (%) of NH_4_^+^ and total nitrogen are defined as the difference between their influent and effluent concentrations divided by the influent concentrations. The influent and effluent were sampled once per week (Figure S3).

During aeration, the aerobic volumetric N-removal rate (mg_N_·L^−1^·d^−1^) was calculated as the difference between the NH_4_^+^ consumption rate and the rates of NO_2_^−^ and NO_3_^−^ production. The aerobic N-removal efficiency (%) was estimated by dividing the N-removal rate during aeration by the NH_4_^+^ depletion rate.

### 2.7 Growth rate of AOB, NOB, and AMX

The maximum growth rates of AOB (μ_AOB,max_) and NOB (μ_NOB,max_) were estimated during *Phase II*, when substrate limitations were minor, based on the measured exponential increase in their maximum activity in the flocs (r_i,max_, Figure 2b), or in their activity during operation (r_i,cycle_, Figure 2c). Most of the activity increase occurred in suspension, where diffusion limitation was assumed to be of minor importance. The suspended solids mass balance (X_i_, with i=AOB, NOB) is expressed as:

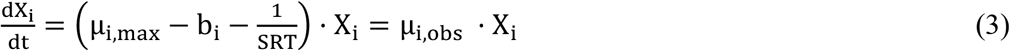

where μ_i,max_ and μ_i,obs_ are the maximum and observed growth rates, respectively, of the guild i (d^−1^), bi is the decay rate of the guild i (d^−1^; set to 0.05 μ_i,max_), and SRT is the solids retention time (d). The value of μ_i,obs_ was obtained from the exponential interpolation of the measured increase in activities (ri, mg_N_·L^−1^·d^−1^):

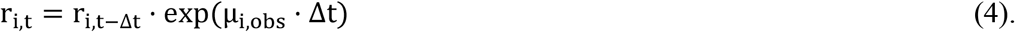

From Eq. 3 and 4, and considering that growth occurs only during the aerobic time, the maximum growth rate can be estimated as follows:

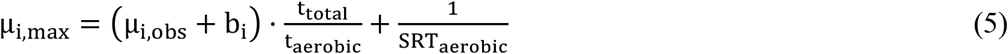

where t_aerobic_/t_total_ is the average fraction of aerobic time over the total batch time, and SRT_aerobic_ the average aerobic SRT during the considered period. The SRT was not considered in the estimation of the maximum growth rate of AMX (μ_AMX,max_), as their growth occurred almost exclusively on the biofilm.

### 2.8 Amplicon sequencing analyses of the bacterial community compositions in biofilm and flocs

The amplicon sequencing method is presented in the Supporting Information, Section S1 (Laureni *et al*., 2016).

### 2.9 Analytical methods

The concentration of NH_4_^+^ was analysed using a flow injection analyser (FIAstar 5000, Foss, Denmark). The concentrations of NO_2_^−^ and NO_3_^−^ were analysed by ion chromatography (Compact IC 761, Metrohm, Switzerland). The COD was measured photometrically with test kits (Hach Lange, Germany). The samples were filtered using 0.45 μm filters (Macherey-Nagel, Germany) prior to analysis. The concentration of total and volatile suspended solids (VSS, TSS) in the mixed liquors was determined according to standard methods (APHA 2005). The total solids (TS) on biofilm carriers were estimated as described previously (Laureni *et al*., 2016).

## 3 Mathematical model of the hybrid system

### 3.1 Model description

A dynamic model of the hybrid MBBR operated in SBR mode was developed and implemented in MATLAB (version R2015b, MathWorks Inc.). The MATLAB scripts are available as open-source code in the Supporting Information. The aim of the model was to understand how the composition of the flocs and the NOB concentration respond to changes in DO, fraction of flocs removed per SBR cycle (f_WAS_), and maximum volumetric AMX activity (r_AMX,max_). To this end, perfect biomass segregation was assumed, with AOB and NOB in the flocs and AMX in the biofilm (Figure 1).

**Figure 1.**
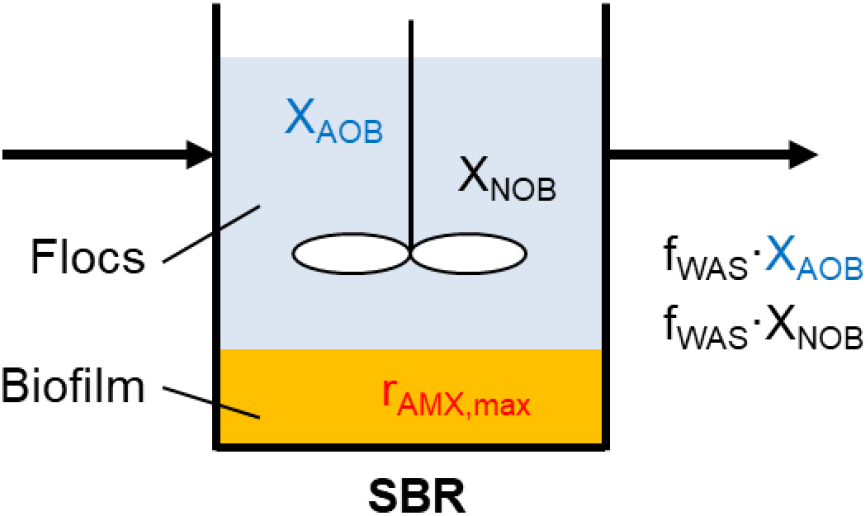
Location of the active biomass in the mathematical model of the hybrid system. The model assumes perfect biomass segregation, with AOB and NOB in the flocs and AMX in the biofilm. r_AMX,max_ is the maximum volumetric anammox activity (mg_(NH4+NO2)-N_·L’ ^−1^·d^−1^). f_WAS_ represents the fraction of flocs removed at the end of each SBR cycle.

Five soluble compounds were considered: ammonium (NH_4_^+^), nitrite (NO_2_^−^), nitrate (NO_3_^−^), di-nitrogen gas (N_2_), and DO.

The AOB, NOB, and AMX processes were modelled according to the stoichiometric and kinetic matrix in Table 1. Unless explicitly stated, parameter values were taken from the literature (Table 2). X_AOB_ and X_NOB_ were assumed to grow in the flocs, and their abundance and activity to be influenced by growth and washout. For the sake of simplicity, the model excluded decay processes. Free ammonia and free nitrous acid inhibitions were considered negligible under mainstream concentrations and pH.

AMX were considered to grow in a deep biofilm (Morgenroth 2008). The primary goal of the modelling was to understand the role of the biofilm as “NO_2_-sink”: the biofilm was consequently modelled as zero-dimensional, and spatial gradients were neglected. In order to discuss the potential effects of diffusion, additional simulations were run with 10-fold increased values for NO_2_^−^ and NH_4_^+^ affinity constants of AMX. Moreover, as the activity of deep biofilms is transport-limited rather than biomass-limited, the maximum AMX process rate (ρ_AMX,max_ = μ_AMX,max_ X_AMX_, mg_COD_·L^−1^·d^−1^; Table 1) was assumed to be constant during each simulation. This was implemented by considering the concentration of AMX (X_AMX_) and the process rate as constants. The oxygen inhibition of AMX was not explicitly modelled: deep biofilms are in fact oxygen-limited, and the modelled AMX activity is to be considered the activity resulting from the anoxic biofilm layers. For consistency with the experimental part, the simulation results are presented as a function of r_AMX,max_ (mg_(NH4+NO2)-N_· L^−1^ · d^−1^) as obtained by the product of ρAMX,max and the sum of the stoichiometric coefficients for NH_4_^+^ and NO_2_^−^ (Table 1).

### 3.2 Simulation strategy and scenario analysis

The influent was assumed to contain 20 mg_NH4-N_·L^−1^ and be devoid of NO_2_^−^, NO_3_^−^, and COD. Filling, settling, and decanting steps were assumed to be instantaneous. Only the aerated phase was simulated dynamically. As in the operation of the experimental reactor, settling was initiated each time the NH_4_^+^ concentration equalled 2 mg_N_·L^−1^; this resulted in variable cycle durations depending on biomass activity. Simulations were performed for a temperature of 15 °C at which maximum growth rates were estimated in the reactor. The DO was assumed constant, and the volumetric exchange of MWW was 50 % per cycle. The initial concentration of NH_4_^+^ at the start of each cycle was the result of mixing (half of its value at the end of the previous cycle plus half of the influent concentration, *i.e*., 11 mg_N_·L^−1^). The NO_2_^−^ and NO_3_^−^ concentrations at the start of each simulated cycle were always equal to half of their values at the end of the previous cycle. A fixed fraction of flocs (f_WAS_) was removed at the end of each cycle. f_WAS_ was defined as the mass removed from the reactor divided by mass of solids present in the reactor, (X_removed_ V_removed_)/(X_reactor_ V_reactor_). Simulations were run until a pseudo steady-state was reached, *i.e*., constant effluent N and flocs concentration. Pseudo steady-state were shown to be independent from the initial X_AOB_ and X_NOB_. The sensitivity of the model outputs was assessed with respect to the ratio between the O_2_ affinity constants of NOB and AOB (K_O2,NOB_/K_O2,AOB_) and the ratio between the NO_2_^−^ affinity constants of NOB and AMX (K_NO2,NOB_/K_NO2,AMX_) (Table S1, Figures S9).

A combination of different ρ_AMX,max_ (0 - 24 mg_COD_·L^−1^·d^−1^; corresponding to r_AMX,max_ 0-300 mg_(NH4+NO2)-N_·L^−1^·d^−1^), and f_WAS_ (0.4 −1.7 %) were simulated for two DO (0.15 and 1.5 mg_O2_·L^−1^). These modelled parameter values were explicitly chosen to fall in the range of the experimental values. To assess the impact of the individual control parameters, four specific scenarios are discussed (Table 3).

**Table 3:** Values of the control parameters for the four tested scenarios.

| Scenario | DO [$\text{mg}_{\text{O}_2}\cdot\text{L}^{-1}$] | $f_{\text{WAS}}$ [%] | $r_{\text{AMX,max}}$ [$\text{mg}_{\text{N}}\cdot\text{L}^{-1}\cdot\text{d}^{-1}$] |
| --- | --- | --- | --- |
| 1 (baseline) | 1.5 | 0.5 | 86 |
| 2 | **0.15** | 0.5 | 86 |
| 3 | 1.5 | **1.7** | 86 |
| 4 | 1.5 | 0.5 | **270** |

### 3.3 Interdependence between f_WAS_, HRT, and SRT

For an SBR where the reaction phase of the cycle is always extended until the target effluent NH_4_^+^ concentration is reached (2 mg_N_·L^−1^), the HRT, the f_WAS_, and ultimately the SRT are interdependent. At pseudo steady-state, the AOB removed at the end of each cycle must equal the growth of AOB during that cycle:

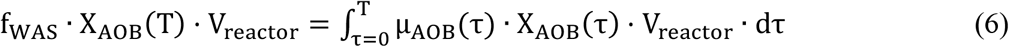

where X_AOB_(T) is the concentration of AOB at the end of a cycle (mg_COD_·L^−1^), T is the length of the cycle (d), V_reactor_ is the working volume of the reactor (L), μ_AOB_(τ) is the actual growth rate of AOB at time τ during the cycle (d^−1^), and X_AOB_(τ) is the AOB concentration at time τ (mg_COD_·L^−1^). Under the simplifying assumption that over a cycle μ_AOB_ ≈ const. and X_AOB_ ≈ const., Eq. 6 can be simplified to

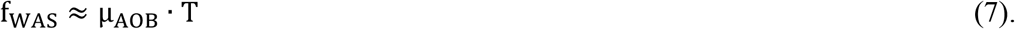

From Eq. 7 it can be seen that the HRT and the cycle time are directly linked: for a given actual growth rate of AOB, increasing f_WAS_ increases T, and thus the HRT. As a result, HRT and f_WAS_ cannot be controlled independently. The value of f_WAS_ also impacts the pseudo steady-state X_AOB_ and X_NOB_, and lower biomass concentrations result from higher f_WAS_. Furthermore, this has direct implications on the SRT of the flocs, defined as the average biomass present in the reactor divided by the biomass removed per cycle. Under the simplifying assumption that X ≈ const. over a cycle, it follows that

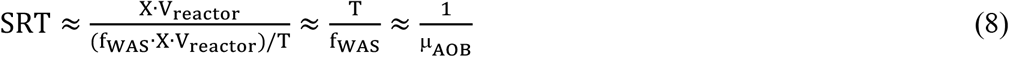

From Eq. 8, after substituting Eq. 7, it can be seen that the SRT is not an independent parameter either, but is directly determined by the actual growth rate of the AOB for the given environmental conditions.

## 4 Results and Discussion

### 4.1 Long term operation of the hybrid MBBR, and the impact of DO on NOB control

#### 4.1.1 Maximum volumetric activities (r_i,max_) segregation between biofilm and flocs

A 12-L hybrid MBBR was operated for mainstream PN/A at 15 °C on aerobically pre-treated MWW, and the impact of the DO on microbial competition and NOB control was investigated. The total and flocs-associated maximum volumetric activities (ri,max) of the three main guilds were measured as proxy for their abundance (Figures 2a, b).

**Figure 2.**
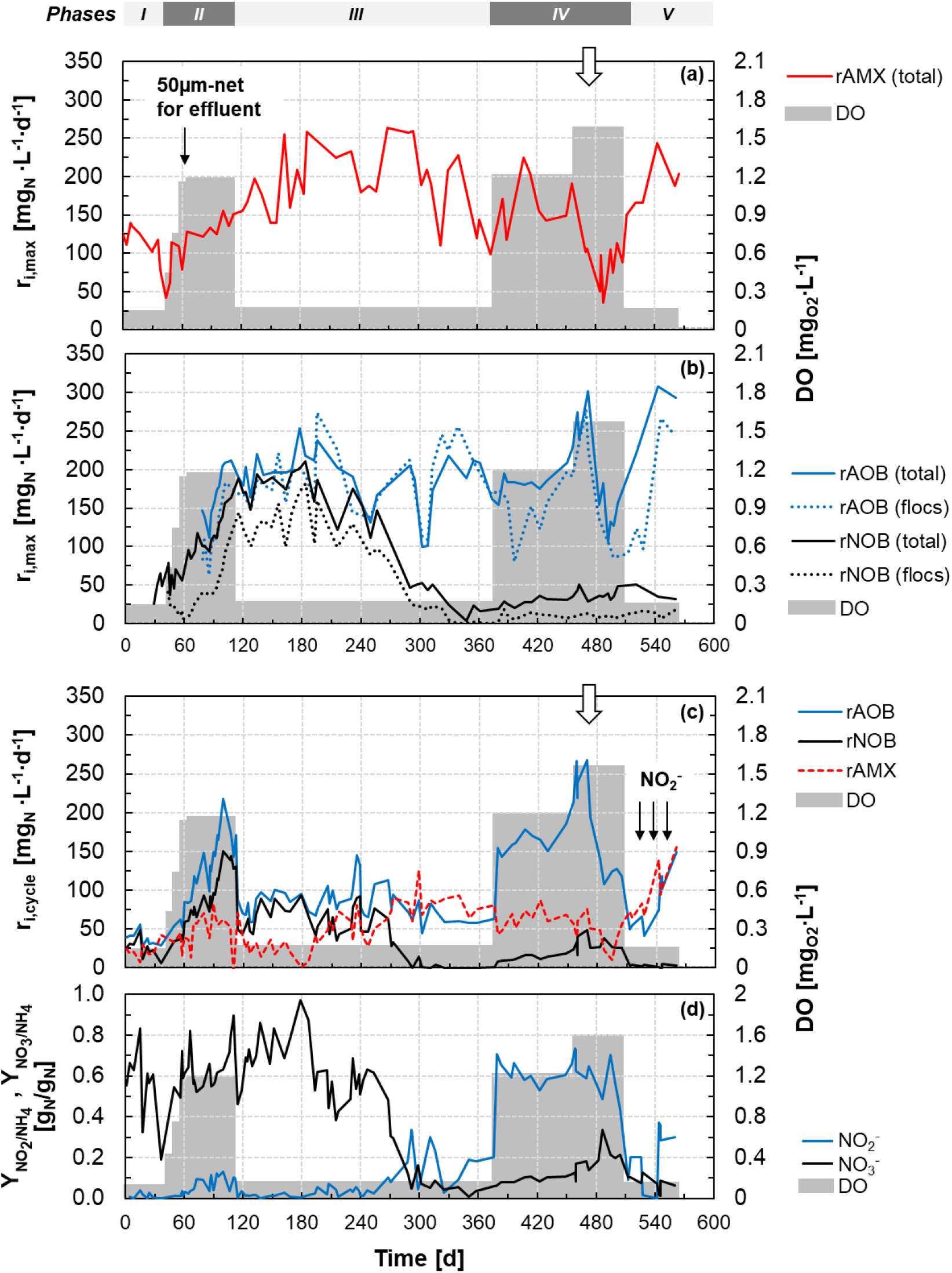
Time series of the maximum (r_i,max_) and actual (r_i,cycle_) volumetric activities of AOB, NOB, and AMX in the hybrid MBBR. (**a**) Total maximum volumetric activities of AMX (the activity in the flocs was negligible throughout the experimental period). (**b**) Segregation of maximum volumetric activities of AOB and NOB: total biomass (biofilm and flocs) and floc fraction only. (**c**) Actual volumetric activities measured during the aerobic phase of an SBR cycle. Activities are expressed as follows: AOB, mg_NH4-N_·L^−1^·d^−1^; NOB, mg_NO3-N_·L^−1^·d^−1^; AMX, mg_(NH4+NO2)-N_·L^−1^·d^−1^. (**d**) Yields of NO_2_^−^ and NO_3_^−^ accumulated relative to the NH_4_^+^ consumed during the aerobic phase. *Shaded area:* the average of the DO concentration measured during aeration over the representative periods. *Vertical black arrows:* in (**a**) time when floc retention was improved by filtering the effluent through a 50-μm-mesh sock-net; in (**c**) time when the volumetric activities during regular operation were measured under non-limiting nitrite concentrations. *Vertical empty arrows:* in (**a, c**) time of the prolonged rain event.

Over more than one year the reactor was stably operated as PN/A (*i.e*. prior to *Phase I* in Fig. 2; (Laureni *et al*., 2016)). During *Phase II*, as a result of the simultaneous increase in DO from 0.17 to 1.2 mg_O2_·L^−1^ and the improved flocs retention, r_AOB,max_ and r_NOB,max_ increased exponentially (Figure 2b). The observed increase was mainly associated with the flocs (dotted line in Figure 2b). Over the same period, the total suspended solids increased from 0.2 to 1 g_TSS_·L^−1^ (Figure S2). The estimated maximum growth rate of AOB (μ_AOB,max_) and NOB (μ_NOB,max_) were 0.30 and 0.34 d^−1^, respectively. For AMX, a μ_AMX,max_ of 0.017 d^−1^ was estimated.

The increase in r_AOB,max_ and r_NOB,max_ stopped when the DO was decreased to its initial value of 0.17 mg_O2_·L^−1^ (day 115, *Phase III*) while keeping all other operational conditions unchanged. After an apparent delay of over six weeks, r_NOB,max_ started to decrease while the established r_AOB,max_ was maintained in the system (Figure 2b). The loss in r_NOB,max_ was primarily associated with the flocs.

During *Phase IV*, r_AOB,max_ and r_NOB,max_ increased exponentially, in particular when the DO was increased to 1.6 mg_O2_·L^−1^ (day 460). Unfortunately, the increase stopped on day 475, when a dramatic drop in all ri,max was observed in correlation with a multiple-day heavy rain event. This also coincided with a 15% loss of TSS in the system, although this alone cannot explain the activity loss. Importantly, all ri,max naturally recovered in less than two months (*Phase V*, Figure 2). All operational conditions are presented in Figure S1.

#### 4.1.2 Volumetric activities during regular operation (r_i,cycle_)

The actual volumetric activities (r_i,cycle_) of the three main guilds were measured during the aerobic step of an SBR cycle to assess the impact of the imposed operational condition on microbial competition. Actual activities are presented in Figure 2c, and the observed yields of NH_4_^+^ converted to NO_2_^−^ and NO_3_^−^ are displayed in Figure 2d.

During periods of high DO (*Phase II and IV*), the volumetric activities during regular operation (r_i,cycle_) were comparable to the maximum activities (ri,max), indicating that substrate limitations were minor under these conditions (Figures 2a, c). The μ_AOB,max_ (0.273 d^−1^) and μ_NOB,max_ (0.286 d^−1^), estimated during *Phase II*, were in good agreement with those obtained from the increase in r_i,max_.

Decreasing the DO on day 115 (*Phase III*) resulted in an immediate decrease of r_AOB,cycle_ and r_NOB,cycle_, as both guilds become DO limited (Figure 2c). After a delay of about two months, r_NOB,cycle_ started to decrease progressively in accordance with the behaviour of r_NOB,max_. The decrease in r_NOB,cycle_ coincided with the increase of r_AMX,cycle_, indicating a progressive shift in the competition for NO_2_^−^. From day 285 onwards, very little NOB activity was detected as supported by the low NO_3_^−^ production. The slight NO_2_^−^ accumulation indicated an excess of r_AOB,cycle_ over the available r_AMX,cycle_ (Figure 2d).

The increase in DO on day 375 (*Phase IV*) led to a sharp increase in r_AOB,cycle_ and lead, due to the excess AOB maintained in the system, to a pronounced accumulation of NO_2_^−^ to about 60% of the consumed NH_4_^+^ (Figure 2d). The rNOB,cycle also increased immediately, due to the NOB persisting in the biofilm, and NO_3_^−^ started to accumulate. The exponential increase of r_AOB,cycle_ and r_NOB,cycle_ stopped on day 475 in conjunction with the heavy rain event (Figure 2c, empty arrow).

#### 4.1.3 Bacterial community composition of biofilm and flocs

The relative read abundances of AOB, NOB, and AMX in the biofilm and flocs are presented in Figure 3. The dynamics of all individual OTUs detected within the three guilds are shown in Figure S4. In good agreement with the observed r_AMX,max_, AMX were almost exclusively present in the biofilm with relative abundances of up to 15% of the total reads (< 0.1% in suspension). Interestingly, four different OTUs were detected for AMX in the biofilm and displayed different dynamics, suggesting possible fine-scale differentiation in the *“Ca*. Brocadia” lineage. Fluorescence *in situ* hybridization (FISH) micrographs of biofilm cryosections are shown in Figure S7.

**Figure 3.**
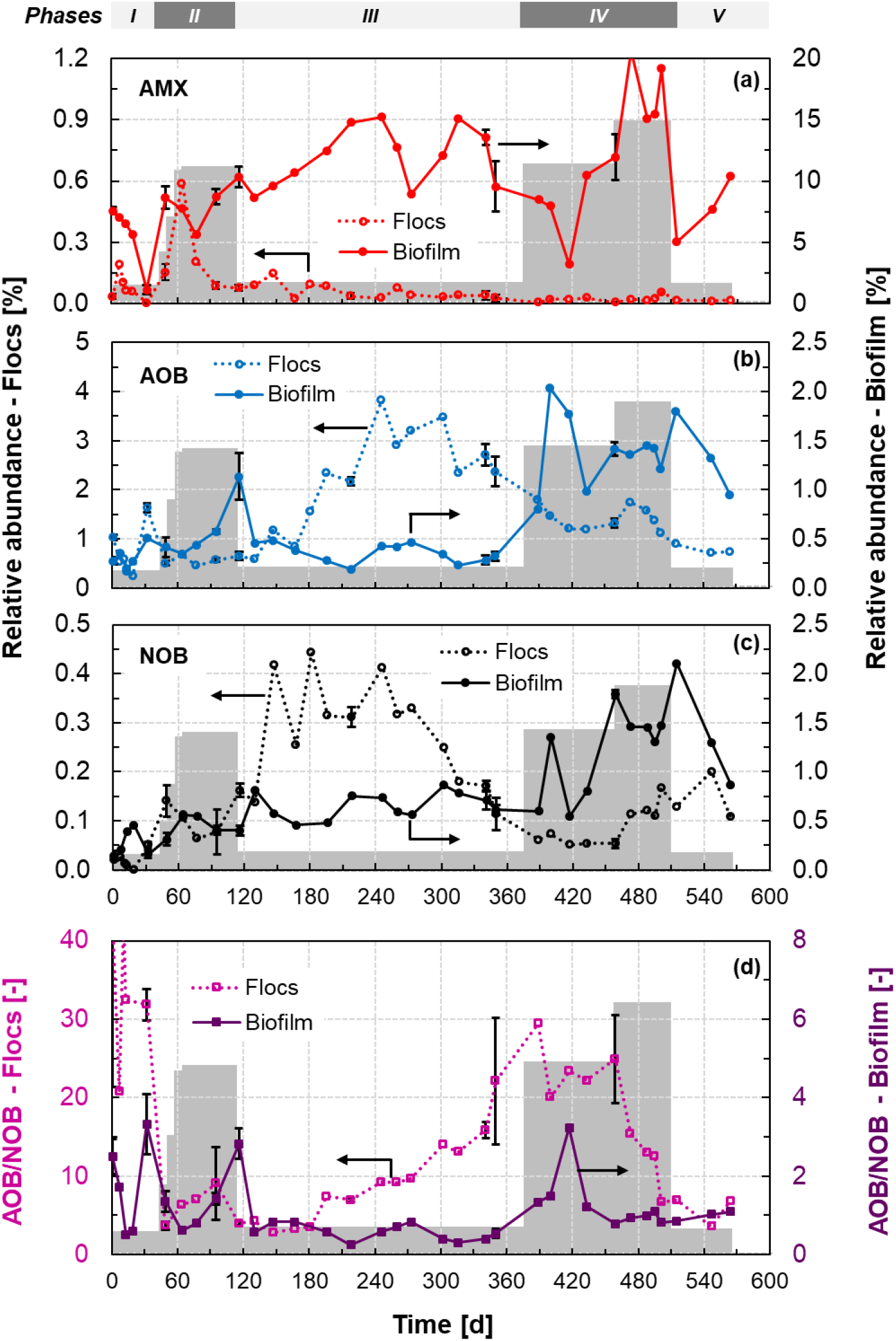
Time series of the relative abundances of AMX (**a**), AOB (**b**), and NOB (**c**) in the flocs (left y-axis) and biofilm (right y-axis) as estimated by 16S rRNA gene-based amplicon sequencing analysis. The displayed values represent the sum of the relative abundances of all OTUs detected for each guild. For the time series of the single OTUs, see Figure S4. (**d**) Time series of the ratio of the relative abundances of AOB and NOB in both the floc and biofilm fractions. *Shaded area*: average operational DO concentration over the representative periods (for values, see Figure 2). *Error bars:* standard deviation of biological triplicates.

Significantly lower relative read abundances were observed for AOB and NOB throughout the entire operation (Figures 3b, c). During *Phase III*, the TSS increased from 1 to over 2.5 g_TSS_·L^−1^ (Figure S2). The relative abundance of AOB (genus *Nitrosomonas*) progressively increased from approximately 0.5 to over 2.5% in the flocs, whereas the relative abundance of NOB (genus *Nitrospira*) decreased progressively from 0.4 to below 0.1%. Thus, the observed loss of NOB activity (Figure 2) coincided with the actual washout of NOB from the flocs. The relative read abundances of both AOB and NOB guilds during *Phase IV* increased markedly on the biofilm, supporting the observed increases in r_AOB,max_ and r_NOB,max_ (Figure 2). Two different OTUs were identified for AOB with distinct trends in biofilm and flocs.

The ratio of the relative read abundances of AOB and NOB is shown in Figure 3d. AOB were selectively enriched over NOB in the flocs during the period at low DO (*Phase III*); the AOB/NOB ratio increased from 5 to over 20. No major changes in the AOB/NOB ratio were observed in the biofilm.

#### 4.1.4 NOB control at low DO: wash-out from the flocs and activity suppression in the biofilm

AOB and NOB grew in the flocs and biofilm. The enrichment of both guilds in the flocs, less diffusion-limited, is in good agreement with previous experimental and modelling reports on PN/A (Hubaux *et al*., 2015, Park *et al*., 2014, Veuillet *et al*., 2014, Vlaeminck *et al*., 2010, Volcke *et al*., 2012, Winkler *et al*., 2011). Also, AOB and NOB displayed comparable maximum specific growth rates as expected at mainstream temperatures (Hellinga *et al*., 1998). In principle, these conditions would hinder the possibility to differentiate the actual growth rates of the two guilds and selectively wash out NOB as efficiently achieved in sidestream suspended biomass systems (Hellinga *et al*., 1998, Joss *et al*., 2011). Nevertheless, prolonged operation at low DO (0.17 mg_O2_·L^−1^) did result in the selective wash out of NOB from the flocs (Figure 2). This is explained by a distinctive characteristic of hybrid systems, namely the competition for NO_2_^−^ between the NOB in the flocs and the AMX enriched in the biofilm acting as a “NO_2_-sink”. The proposed mechanisms for the selective NOB washout are extensively discussed in the modelling section.

The accumulation and persistence of an NOB fraction in biofilms has also been widely reported, and makes the suppression of NO_2_^−^ oxidation challenging in solely biofilm PN/A systems (Fux *et al*., 2004, Gilbert *et al*., 2015a, Isanta *et al*., 2015, Lotti *et al*., 2014, Park *et al*., 2014, Poot *et al*., 2016, Veuillet *et al*., 2014). Here, the actual nitratation activity of the NOB (r_NOB,cycle_) in the biofilm was consistently controlled by the DO, and was completely suppressed at 0.17 mg_O2_·L^−1^ (*Phase III and V*) presumably due to diffusion limitations. To assess whether rNOB,cycle was suppressed only by DO limitation or also by NO_2_^−^ limitation, ri,cycle were measured under non-limiting NO_2_^−^ concentrations. No increase in rNOB,cycle was observed, confirming that DO rather than NO_2_^−^ was the limiting substrate for NOB in the biofilm (Figure 2c, vertical black arrows in *Phase V*). As a result of the selective enrichment of AOB in the flocs, high NO_2_^−^ fluxes to the biofilm for AMX can be guaranteed at sufficiently low DO to suppress NOB activity in the biofilm.

#### 4.1.5 Effluent quality

Overall, the wash-out of NOB from the flocs and the suppression of their activity in the biofilm at low DO, resulted in N-removals over 88 ± 4% and a residual concentration of total N below 3 mg_N_·L^−1^ (1.9 ± 0.5 mg_NH4-N_·L^−1^, 0.3 ± 0.2 mg_NO2-N_·L^−1^, and 0.5 ± 0.3 mg_NO3-N_·L^−1^). This is the highest effluent quality reported so far for mainstream PN/A systems (De Clippeleir *et al*., 2013, Gilbert *et al*., 2015a, Laureni *et al*., 2016, Lotti *et al*., 2014). Moreover, the aerobic N-removal rates achieved (79 ± 16 mg_N_·L^−1^·d^−1^), at an HRT of 11 ± 2 h, were comparable to those of conventional WWTP (Metcalf & Eddy *et al*., 2013). The dynamics of influent and effluent concentrations are presented in Figure S3.

### 4.2 Mathematical modelling of the hybrid MBBR

A simple dynamic model was developed to understand how the NOB concentration in the flocs (X_NOB_), respond to changes in DO, fraction of flocs removed per SBR cycle (f_WAS_), and maximum volumetric AMX activity in the biofilm (r_AMX,max_). To assess the impact of the individual control parameters four different scenarios were simulated (Table 3). The dynamics of X_AOB_ and X_NOB_, and effluent N concentrations are presented in Figure 4, and one cycle at pseudo steady-state is shown for each scenario in Figure S5. The interdependences between the parameters and the impacts of substrate affinities are also discussed.

**Figure 4.**
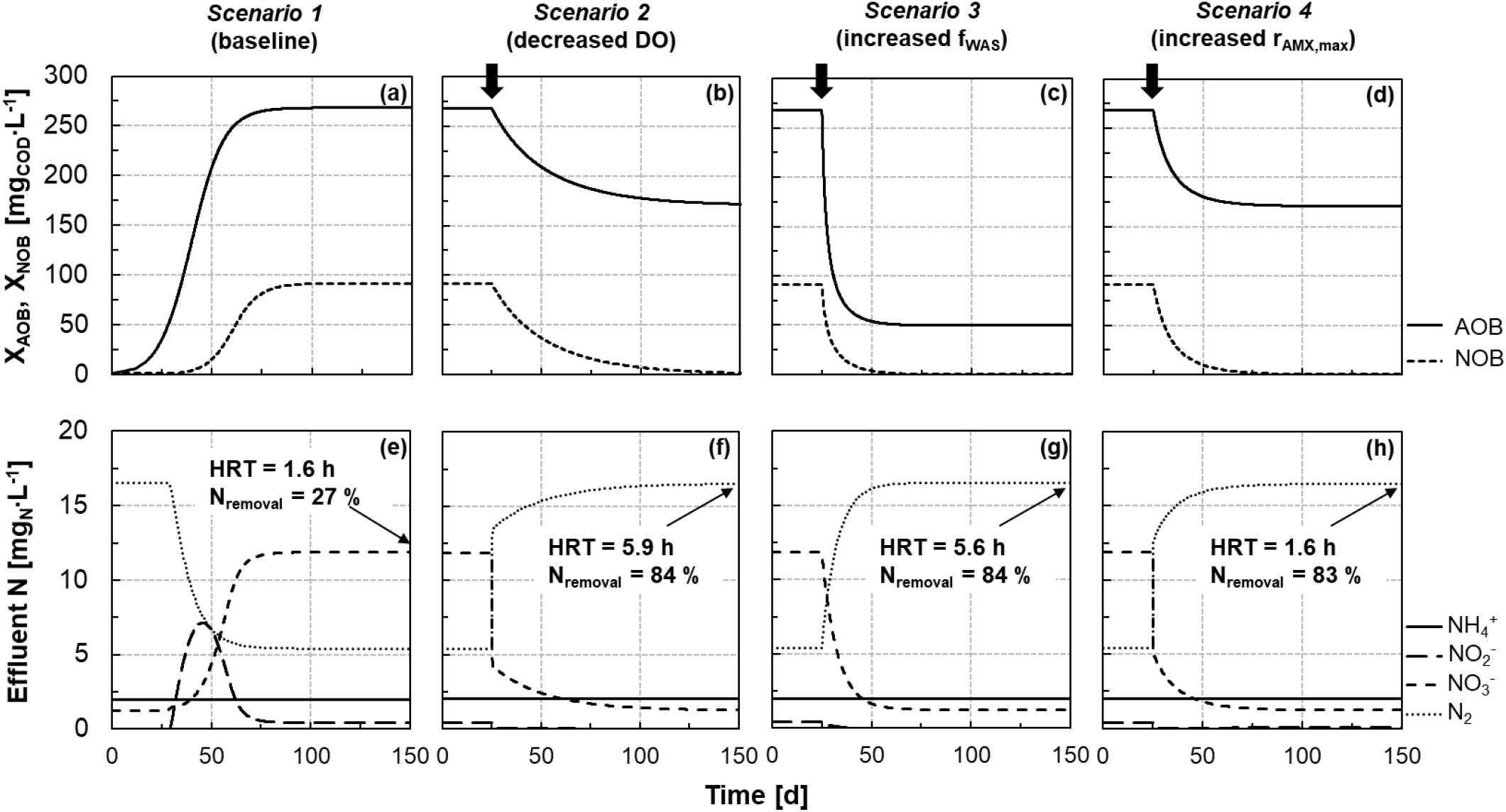
Results from mathematical modelling of dynamics in concentrations of AOB (X_AOB_), NOB (X_NOB_), and effluent N towards the pseudo steady-state for the four scenarios detailed in Table 3. Pseudo steady-state in *Scenario 1* is used as initial conditions for *Scenarios 2, 3*, and *4*. Profiles of nitrogen species and biomass evolution during an SBR cycle at pseudo steady-state for the four scenarios are presented in Figure S6. *Vertical thick arrows:* times when scenario-specific modification of operational conditions was implemented.

#### 4.2.1 Scenario 1 (baseline): high AOB and NOB enrichment in the flocs

A low initial concentration of 1 mg_COD_·L^−1^ was set for X_AOB_ and X_NOB_. Prolonged operation at 1.5 mg_O2_·L^−1^ resulted in the enrichment of both AOB and NOB in the flocs (Figure 4a), similar to experimental observations during reactor operation (*Phase II*, Figure 2). The pseudo steady-state X_AOB_ and X_NOB_ obtained in *Scenario 1* were assumed as initial concentrations for the other scenarios.

#### 4.2.2 Scenario 2: the DO controls the selective washout of NOB from the flocs

The DO has a direct impact on the growth rate of both AOB and NOB (see process rates in Table 1). AOB and NOB are also equally exposed to washout, *e.g*. by removing a fraction of flocs at the end of each SBR cycle (f_WAS_). However, only the NOB growth rate is impacted by the competition for NO_2_^−^ with the “NO_2_-sink” represented by the AMX in the biofilm. This direct competition for NO_2_^−^ between NOB and AMX leads to a difference in the actual growth rates of AOB and NOB (*i.e*., μ_NOB_ < μ_AOB_) providing the basis for the selective NOB washout (*i.e*., μ_NOB_ < SRT^−1^ < μ_AOB_).

The impact of a DO decrease to 0.15 mg_O2_·L^−1^ was assessed in *Scenario 2* to reflect the experimental strategy (*Phase III*, Figure 2). Under the imposed DO-limiting condition, and at the fixed f_WAS_, only AOB could be maintained in the system while NOB were successfully washed out. High N-removals are achieved (84%; Figures 4b, f). At the same time, due to the decreased AOB activity the HRT increases from 1.6 to 5.9 h (i.e. longer cycles are required to achieve the set effluent NH_4_^+^ concentration). In terms of effluent concentrations, the reduction of the DO limits the aerobic activity (as was the case in the reactor, Figure 2c) and results in the immediate reduction of NO_3_^−^ (Figure 4f).

The numerical results provide a mechanistic interpretation for the experimental observations: the sole reduction of the DO was sufficient to reduce the actual NOB growth rate below the minimum required to prevent their washout. Moreover, the simulations support the possibility to use DO to achieve the selective washout of NOB from the flocs.

#### 4.2.3 Scenario 3: increasing the fraction of flocs removed per cycle is an effective strategy to achieve selective NOB washout

Decreasing the DO might not always be a viable option at full scale, either because the operational DO is already low or the size of the installed aerators and blowers is not suitable (Joss *et al*., 2011). Conversely, the selective removal of the flocs from a hybrid MBBR, or of fine particles from a granular sludge system, may be a more feasible option, *e.g*., via a separate settler (Veuillet *et al*., 2014), hydrocyclone (Wett *et al*., 2015), or screen (Han *et al*., 2016). Simulations were run to assess the effectiveness of increasing the fraction of flocs removed at the end of each SBR cycle as a strategy to achieve the selective washout of NOB.

Numerical results suggest that successful NOB washout can indeed be achieved by increasing f_WAS_ while maintaining all other conditions unchanged. Under *Scenario 3*, only the f_WAS_ was increased to 1.7 % and, as a result, NOB were selectively washed out (Figure 4c). In this case, the actual NOB growth rate (function of DO and NO_2_^−^ concentrations, Table 1) is no longer sufficient to compensate for the increased washout. Simultaneously, the significantly lower AOB concentrations maintained in the system result in higher HRT and thus reduced N-loads that can be treated at the same effluent quality (Eq. 7). Nevertheless, in comparison to lowering the DO, increasing f_WAS_ allows a faster NOB washout. From a process control perspective, the proposed simulation examples highlight how in principle NOB can be washed out by only controlling the removal of the flocs.

#### 4.2.4 Scenario 4: variations of AMX activity in the biofilm - the “NO_2_-sink” - have a direct impact on NOB concentration in the flocs

The NOB in the flocs compete for NO_2_^−^ with the AMX enriched in the biofilm - the “NO_2_-sink” - here represented by the maximum volumetric AMX activity (r_AMX,max_). Increasing r_AMX,max_, *i.e*. the rate of NO_2_^−^ consumption by AMX, reduces the bulk NO_2_^−^ concentration and consequently the actual NOB growth rate analogously to decreasing the DO.

The possibility of achieving complete and selective NOB washout from the flocs by increasing r_AMX,max_ was shown numerically. Under *Scenario 4*, the increase in r_AMX,max_ resulted in a higher NO_2_^−^ consumption, and thus a stronger competition with NOB, which are successfully washed out (Figure 4d). At the same time, simulations indicate that increasing r_AMX,max_ results in slightly lower AOB concentrations, as AMX reduce the NH_4_^+^ available for AOB growth, with however minor implications in terms of HRT. As a result, a high N-removal is achieved while still maintaining a low HRT. The dynamics in effluent N concentrations are similar to *Scenario 2*. An immediate decrease of the NO_3_^−^ concentration, due to the reduced NO_2_^−^ available for NOB, is followed by a further progressive reduction as NOB are washed out (Figure 4h).

At full scale, the maximum AMX activity can in principle be increased, *e.g*. by bio-augmentation from a sidestream PN/A process (Wett *et al*., 2015). On the other hand, a partial or complete inhibition of the AMX guild represents the opposite case where NOB may grow in the flocs due to the reduced competition for NO_2_^−^. Under such circumstances, increasing f_WAS_ and/or reducing the DO may be suitable operational strategies to prevent NOB proliferation, as will be discussed in the next section.

#### 4.2.5 Interdependent impacts of DO, f_WAS_, and r_AMX,max_, on NOB, and the impact of substrates diffusion in the biofilm

To better understand the interdependence between the different control parameters, the pseudo steady-state concentrations of X_AOB_, X_NOB_ and effluent NO_3_^−^ are shown in Figure 5 as a function of different r_AMX,max_ and f_WAS_. Two DO concentrations were simulated (0.15 and 1.5 mg_O2_·L^−1^), representative of the low and high DO experimental periods. The pseudo steady-state of the four scenarios discussed in the previous sections are highlighted.

**Figure 5.**
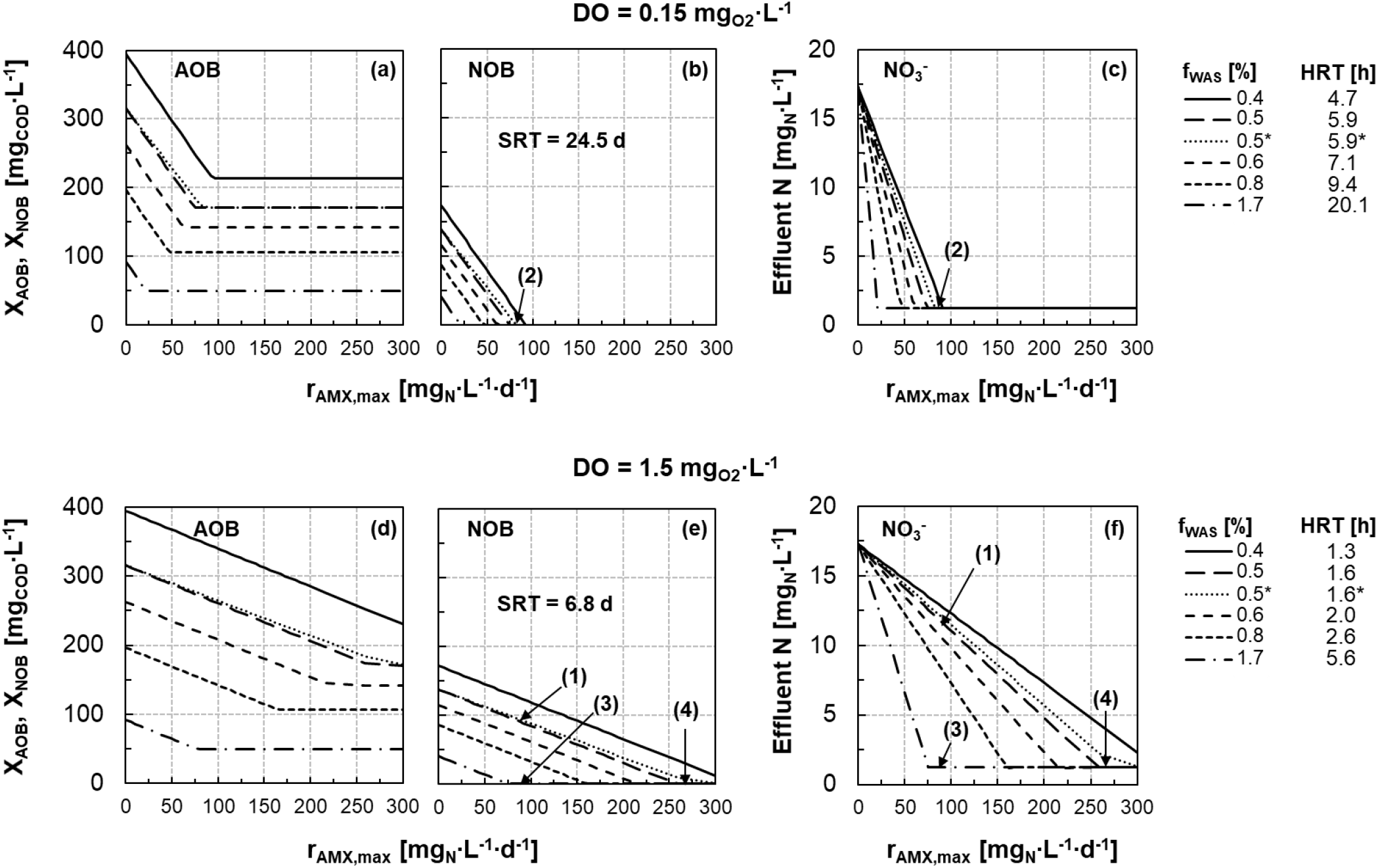
Concentrations of AOB (**a, d**) and NOB (**b, e**) in the flocs under pseudo steady-state conditions modelled as a function of the maximum volumetric AMX activity (r_AMX,max_ mg_(NH4+NO2)-N_·L^−1^·d^−1^) for two reference DO, 0.15 and 1.5 mg_O2_·L^−1^. (**c, f**) Residual concentration of NO_3_^−^ in the effluent at pseudo steady-state. NH_4_^+^, NO_2_^−^ and N2 concentrations are presented in Figure S5. The different lines represent different f_WAS_ values, as shown in the legend to the right of the figures. The resulting HRT for each f_WAS_ is also reported in the legend. Simulations were run with reference parameters shown in Table 2. Only for the case marked with (*), the ammonium and nitrite affinity constants of AMX were increased by a factor of ten. *Black arrows and numbers in parentheses:* the four scenarios discussed in the text and presented in Figure 4.

XNOB and the effluent NO_3_^−^ concentration decrease with increasing ^r^AMX,max (i.e. the competing “NO_2_-sink”). For any given DO and f_WAS_, there is a minimum r_AMX,max_ required for full NOB washout from the flocs (Figures 5b, e). X_AOB_ also decrease with increasing r_AMX,max_. In fact, by consuming NH_4_^+^, AMX reduce its availability for AOB growth (Figures 5a, d). This effect disappears, and X_AOB_ stabilizes, as soon as the NOB are fully washed out. As a matter of fact, when present in the system, NOB consume NO_2_^−^ and indirectly favour AOB by decreasing NH_4_^+^ depletion by AMX. As an example, the case of partial AMX inhibition would be equivalent to moving horizontally to the left in Figure 5: an increased X_NOB_ is to be expected unless *e.g*. DO is decreased or/and f_WAS_ is increased.

Additional simulations with a conservative ten-times higher value for both NH_4_^+^ and NO_2_^−^ affinity constants of AMX were run to assess the effects of substrate diffusion through the biofilm on the modelled pseudo steady-states. Only the case of f_WAS_ equal to 0.5% was considered. As can be seen from Figure 5, differences from the reference case (i.e. with unmodified affinity constants) are negligible. It is therefore deemed justified to neglect diffusion effects for the purpose of this work.

Overall, when interpreting the numerical results, it is important to consider the simplifying assumptions made in the modelling of the biofilm. AMX inhibition by oxygen was neglected, and the r_AMX,max_ was assumed to be the result of the active AMX in the anoxic layers of a deep biofilm. In addition, no NOB growth in the biofilm was considered. In this respect, it is worth noting that the nitrifying activity of NOB was shown experimentally to be completely suppressed at low DO. Additional simulations with more complex models, including biomass stratification and inhibition processes, are recommended here. Nevertheless, the simplified model allowed to identify the fundamental role played by the AMX-enriched biofilm (“NO_2_-sink”) in favouring the selective NOB washout from the flocs.

#### 4.2.6 The possibility of successful NOB washout from the flocs is not impaired by the values of the affinity constants

In solely biofilm PN/A systems, the ratio of the oxygen affinity constants, K_O2,NOB_/K_O2,AOB_, and the ratio of the NO_2_^−^ affinity constants, K_NO2,NOB_/K_NO2,AMX_, are reported as the main parameters controlling microbial competition (Brockmann and Morgenroth 2010, Hao *et al*., 2002, Pérez *et al*., 2014, Picioreanu *et al*., 2016). For example, Hao *et al*., (2002) have reported that K_O2,NOB_/K_O2,AOB_ > 0.2 and K_NO2,NOB_/K_NO2,AMX_ > 3 is a required condition for successful NOB suppression in a biofilm system modelled at 30°C. In the present study, the sensitivity of the simulation results and the validity of the previously drawn conclusions was tested with respect to the ratios K_O2,NOB_/K_O2,AOB_ and K_NO2,NOB_/K_NO2,AMX_. To ease the interpretation of the sensitivity analysis, K_O2,AOB_ was maintained constant (0.6 mg_O2_·L^−1^), and the K_O2,NOB_/K_O2,AOB_ ratio was varied between 0.14 (Regmi *et al*., 2014) and 2.00 (Perez *et al*., 2014) by changing K_O2,NOB_ (Table S1). Simulations were run for the two reference DO of 0.15 and 1.5 mg_O2_·L^−1^, and a fixed f_WAS_ of 0.5%. The pseudo steady-state X_NOB_ and effluent NO_2_^−^ concentrations are displayed as a function of K_O2,NOB_/K_O2,AOB_ in Figure 6. An overview of X_AOB_ and X_NOB_, and the effluent concentrations of the dissolved N species, is presented in Figure S8.

**Figure 6.**
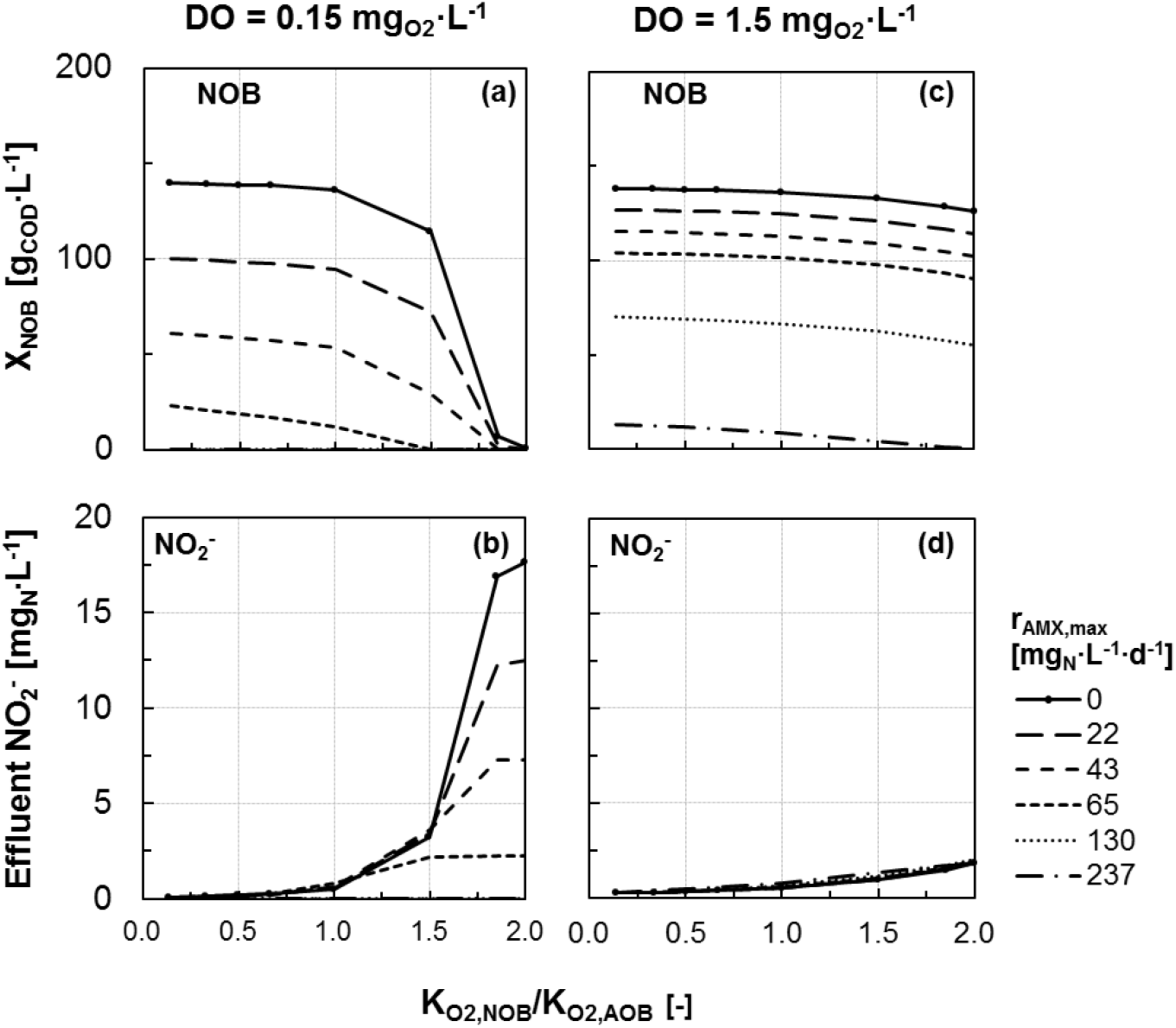
Sensitivity analysis. Impact of different K_O2,NOB_/K_O2,AOB_ on simulated NOB concentrations at pseudo steady-state (**a, c**) and corresponding effluent NO_2_^−^ concentrations (**b, d**) for the two reference DO (0.15 and 1.5 mg_O2_·L^−1^). K_O2,NOB_/K_O2,AOB_ = 0.67 is the reference case (see Table 2). The values of the oxygen affinities for NOB and AOB and their ratio are presented in Table S1. In the simulations, an f_WAS_ of 0.5% was assumed. All concentrations of X_AOB_ and effluent N species at pseudo steady-state are presented in Figure S8. r_AMX,max_ is expressed as mg_(NH4+NO2)-N_·L^−1^ ·d^−1^.

At a low DO (0.15 mg_O2_·L^−1^), the value of K_O2,NOB_/K_O2,AOB_ determines the mechanisms controlling NOB washout. On the one hand, for values of K_O2,NOB_/K_O2,AOB_ < 1, low NO_2_^−^ concentrations are modelled (i.e. rapidly consumed by NOB and AMX), and the competition with AMX for NO_2_^−^ is the dominant mechanism controlling NOB washout. Increasing r_AMX,max_ results in lower NOB pseudo steady-state concentrations (Figure 6a). Importantly, NOB are successfully washed out in the model even in the extreme case of K_O2,NOB_/K_O2,AOB_ = 0.14 (Regmi *et al*., 2014), which would make their control challenging in solely biofilm systems (Brockmann and Morgenroth 2010, Hao *et al*., 2002, Pérez *et al*., 2014). On the other hand, for higher values (K_O2,NOB_/K_O2,AOB_ > 1), DO limitation starts to play an important role. Due to the reduced NOB growth rate, lower NOB concentrations can be sustained in the system, and NO_2_^−^ accumulates if the AMX activity is not sufficiently high (Figure 6b).

Interestingly, for large K_O2,NOB_ (K_O2,NOB_/K_O2,AOB_ = 2.00), NOB are washed out from the system even in the absence of AMX and despite high NO_2_^−^ accumulation. In this case, the actual NOB growth rate is not sufficient to maintain them in the system at the cycle length set by AOB and the imposed f_WAS_ (Eq. 7). Importantly, if r_AMX,max_ is sufficiently high (e.g. > 65 mg_N_·L^−1^·d^−1^), the NOB washout does not depend on K_O2,NOB_/K_O2,AOB_.

At a high DO (1.5 mg_O2_·L^−1^), NOB washout is less sensitive to the value of K_O2,NOB_/K_O2,AOB_, and the competition for NO_2_^−^ with AMX is the dominant mechanism controlling NOB washout (Figure 6c). Nevertheless, in analogy to the low DO case, NO_2_^−^ accumulation occurs for high values of K_O2,NOB_/K_O2,AOB_. Taken together, these results provide a mechanistic hypothesis to explain the seemingly contradictory experimental observations during *Phase IV* (Figure 2), when only limited NOB enrichment was observed in the flocs despite high DO and pronounced NO_2_^−^ accumulation. In general, higher r_AMX,max_ are required for NOB washout (e.g., > 237 mg_N_·L^−1^·d^−1^) compared to the case at low DO.

In terms of NO_2_^−^ affinity constants, K_NO2,NOB_ was decreased from a usually assumed value 100 times higher than K_NO2,AMX_ (Hao *et al*., 2002, Pérez *et al*., 2014) to a value of 0.1 K_NO2,AMX_ (Figure S9). Decreasing K_NO2,NOB_ increases the competitive advantage of NOB over AMX and results in higher XNOB at pseudo steady-state for any given r_AMX,max_. Nevertheless, within the broad range of values tested, NOB washout can always be achieved provided that a sufficiently high r_AMX,max_ is present (Figure S9).

In summary, this work strongly support the increased operational flexibility offered by hybrid systems, as compared to solely biofilm systems, for the control of NOB under mainstream conditions. In fact, irrespective of the values chosen for the affinity constants, it is in principle always possible to control the selective pressure on NOB via DO, f_WAS_, and/or r_AMX,max_, and achieve their complete washout.

## 5 Conclusions

This study aimed at understanding the mechanisms underlying microbial competition and the control of NOB in hybrid PN/A reactors. To this end, a hybrid MBBR was operated under mainstream conditions and a simple mathematical model of the system was developed. Experimentally, AMX were shown to enrich in the biofilm while AOB and NOB grew preferentially in the flocs. AMX are retained in the biofilm independent of floc removal and they act as a “NO_2_-sink”. Conversely, AOB and NOB are maintained in the flocs only if their actual growth rates is larger than the imposed washout (*i.e*., if μ > SRT^−1^).

- The key mechanisms for selectively washing out NOB from the system are maintaining a sufficiently low SRT for the flocs and limiting NO_2_^−^ bulk phase concentrations by means of the AMX “NO_2_-sink”. AOB growth rates are not affected by NO_2_^−^ bulk phase concentrations allowing reactor operation with selective washout of NOB while keeping AOB.
- Experimental results and numerical simulations showed that, for an imposed fraction of flocs removed per SBR cycle or given SRT, NOB can be selectively washed out by decreasing the DO-setpoint, *e.g*., from 1.2 to 0.17 mg_O2_·L^−1^. In this case, while both AOB and NOB actual growth rates decrease; due to the concurrent NO2-limitation only NOB growth rate is reduced below the washout threshold *i.e*., μ_NOB_< SRT^−1^ < μ_AOB_.
- In analogy, for a given DO-setpoint, simulations indicated that selective NOB washout can be achieved also by increasing the fraction of flocs removed: the actual NOB growth rate remains unaffected but is no longer sufficient to compensate for the increased washout.
- Moreover, differently from pure biofilm systems where NOB suppression relies on a larger oxygen affinity of AOB than NOB, modelling results suggest that it is in principle always possible to selectively wash out NOB by controlling the DO-setpoint and/or the flocs removal provided AMX act as “NO_2_-sink” in the biofilm.

Ultimately, this study demonstrates the high operational flexibility, in terms of variables that can be easily controlled by operators, offered by hybrid systems for the control of NOB in mainstream PN/A applications.

## Supporting information

## 6 Acknowledgements

This study was funded by the European Research Council ERC via the ATHENE project (grant agreement 267897). ML was partially supported by a Marie Sklodowska-Curie Individual Fellowship (MixAmox project; grant agreement 752992). We sincerely thank Kai Udert and Nicolas Derlon for valuable discussions, Marco Kipf for his support in the laboratory, Brian Sinnet for the particle size analysis, and Claudia Baenninger-Werffeli, Sylvia Richter, and Karin Rottermann for their assistance with the physicochemical analyses of all the samples.

