## Supplementary material for "Biomass segregation between biofilm and flocs improves the control of nitrite-oxidizing bacteria in mainstream partial nitritation and anammox processes"

#### S1 – Amplicon sequencing analysis of the bacterial community compositions in biofilm and flocs

The biofilm and flocs fractions were sampled from the hybrid MBBR on a weekly or bi-weekly basis, and stored at -20°C. Their bacterial community compositions were analysed by sequencing the V4 region of the 16S rRNA gene using high throughput amplicon sequencing as described elsewhere (Caporaso *et al.*, 2012). The biofilm carriers were cut into pieces and inserted directly into lysing matrix E tubes for DNA extraction using Fast DNA Spin Kits for Soil (MP Biomedicals, USA), which were then treated in accordance with the manufacturer's recommendations, with an amendment to the bead beating step to 4 cycles of 40 seconds at 6 m/s. Samples were prepared in triplicate to ensure comparability. Primers 515F (5'-GTGCCAGCMGCCGCGGTAA-3') and 806R (5'-GGACTACHVGGGTWTCTAAT-3') were selected for their known ability to detect AMX and for their broad coverage of bacterial diversity (Gilbert *et al.*, 2014; Weissbrodt *et al.*, 2015), as also applied in (Laureni *et al.*, 2015). An equimolar pool of amplicons was sequenced using a MiSeq sequencer (Illumina, USA) at an average depth of 36,500 ± 4,100 reads (min = 25,476, max = 41,844) per sample. The datasets were processed as described elsewhere (Karst *et al.*, 2016). Operational taxonomic units (OTUs) were clustered at 97% sequence similarity, and taxonomy was assigned using RDP as implemented in QIIME, using MiDAS (v1.20) as a reference database (McIlroy *et al.*, 2015). Known populations of AOB, NOB, and AMX guilds were identified, and their relative abundances assessed. Several populations harboured different OTUs (Figure S4).

**Table S1**

Sensitivity analysis of the model towards changes in $K_{O2,NOB}$/ $K_{O2,AOB}$ (Figure 6): tested values of oxygen affinities constants for NOB and AOB, and the corresponding ratio.

| $K_{O2,NOB}$ $mg_{COD}\cdot L^{-1}$ | $K_{O2,AOB}$ $mg_{COD}\cdot L^{-1}$ | $K_{O2,NOB}$ / $K_{O2,AOB}$ - |
|---|---|---|
| 0.08 | 0.6 | 0.14 |
| 0.2 | 0.6 | 0.33 |
| 0.3 | 0.6 | 0.50 |
| 0.4* | 0.6* | 0.67 |
| 0.6 | 0.6 | 1.00 |
| 0.9 | 0.6 | 1.50 |
| 1.11 | 0.6 | 1.85 |
| 1.2 | 0.6 | 2.00 |

*Reference scenario (Table 2).

### Figures S1-S9

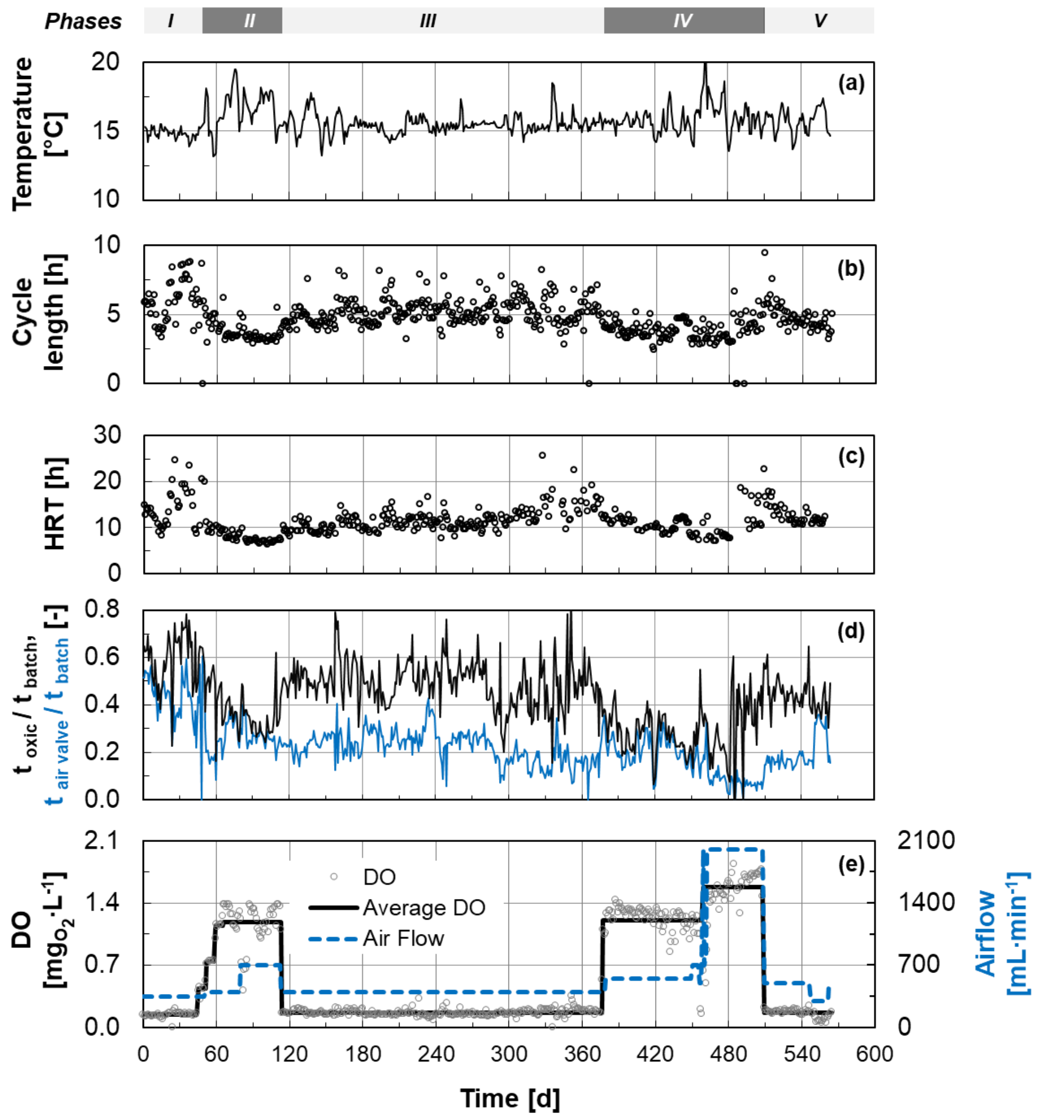


**Figure S1:** Operational conditions of the hybrid MBBR during the whole experimental period. Time series of (**a**) temperature; (**b**) cycle length; (**c**) hydraulic retention time (HRT); (**d**) oxic time over total batch time ($t_{oxic}$ /$t_{batch}$) and time the air valve was open over total batch time ($t_{air\ valve}$/$t_{batch}$); (**e**) daily average dissolved oxygen (DO) concentration (circles), average DO during the different representative periods (continuous black line, also reported in Figures 2, 3 in the main manuscript), and airflow (dashed blue line).

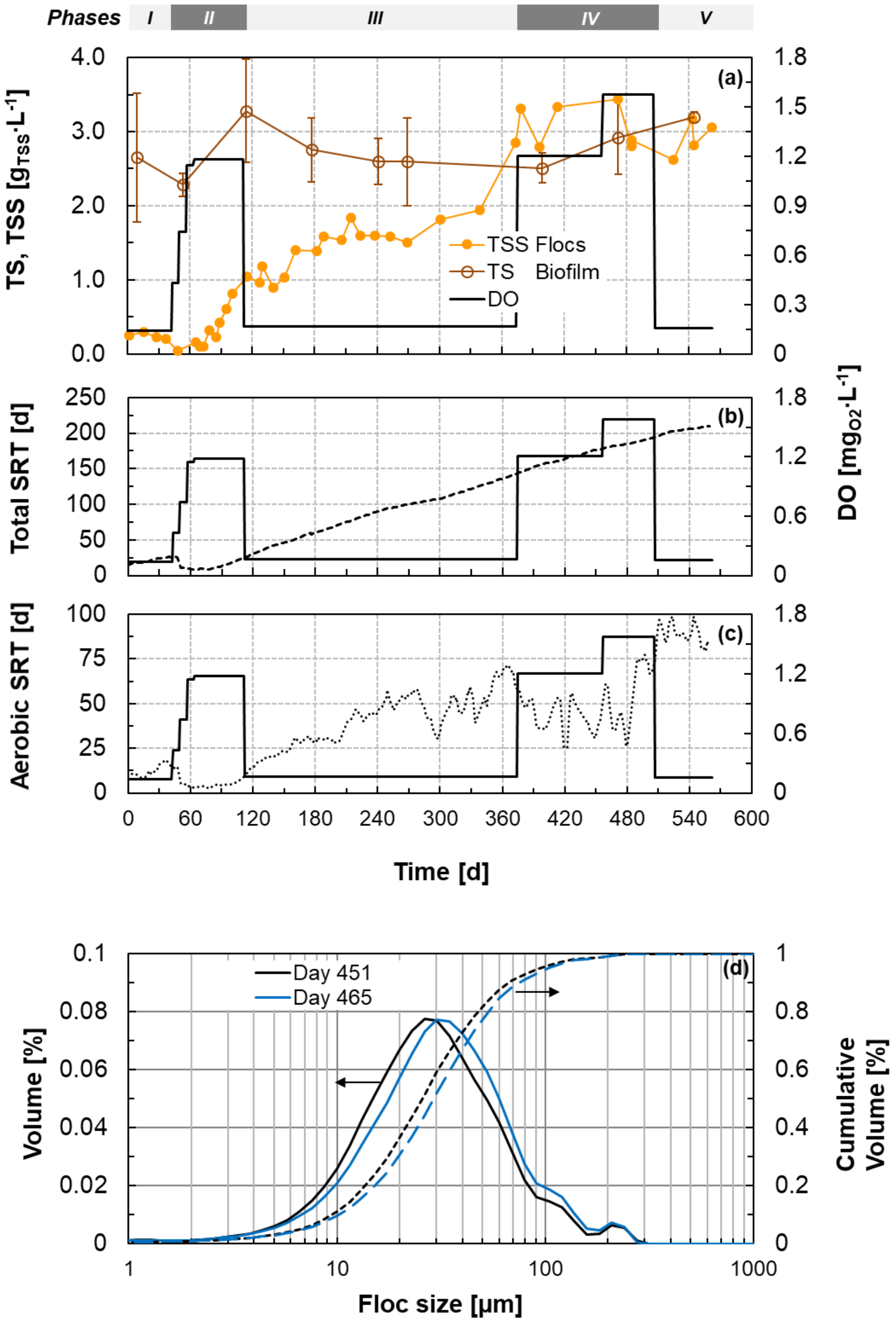


**Figure S2:** (**a**) Temporal evolution of the total solids (TS) of the biofilms attached to the carriers and the total suspended solids of the flocs (TSS). The TSS of the flocs were measured during the second cycle after effluent biomass reintroduction. Error bars for the biofilm fraction represent the standard deviation of three carriers. (**b**, **c**) Temporal evolution of the total and aerobic SRT of the flocs biomass fraction. The higher fluctuations in the aerobic SRT are due to the variability of the aerobic over total time (Figure S1). (**d**) Volumetric particle size distribution of the flocs, measured on days 451 and 465 via laser light scattering, as fraction and cumulative volumes. *Black continuous line* (**a-c**): average DO during the different representative periods, values on right axis.

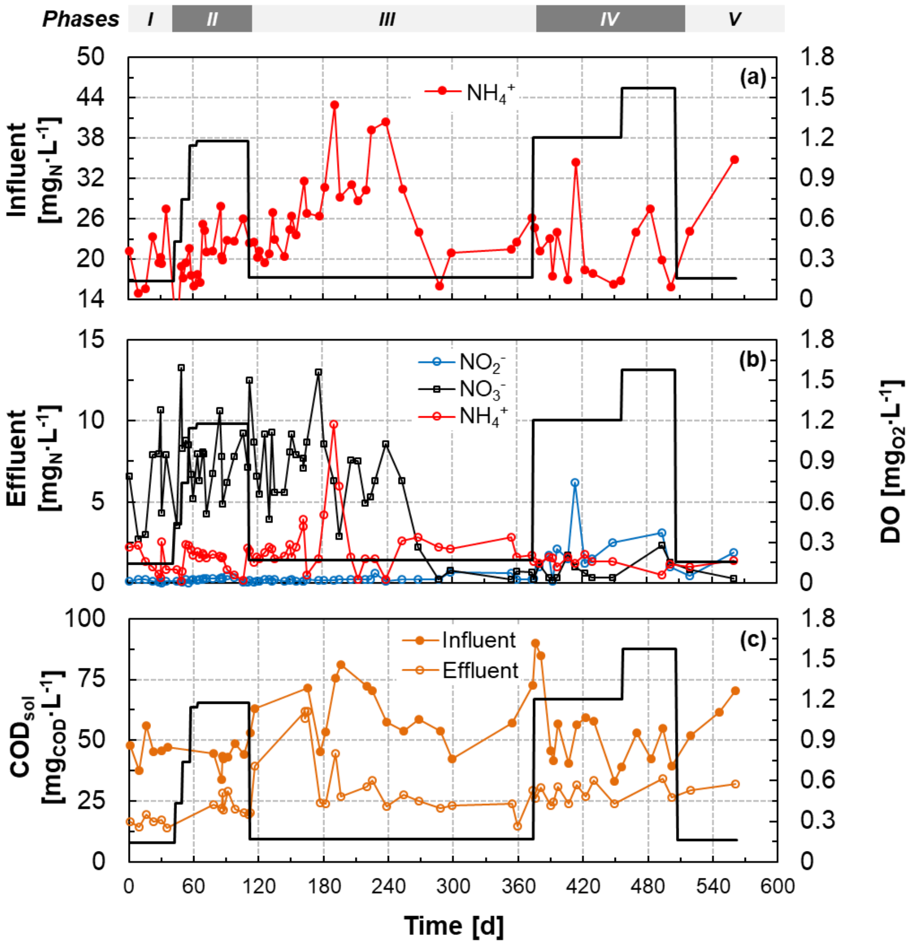


**Figure S3:** Trend of (**a**) $NH_4^+$ concentration in the influent of the hybrid MBBR ($NO_2^-$ and $NO_3^-$ were below the detection limit of 0.2 $mg_N \cdot L^{-1}$ in the majority of samples); (**b**) concentrations of nitrogen species ($NH_4^+$, $NO_2^-$, $NO_3^-$) in the effluent; (**c**) concentration of soluble organic matter (measured as chemical oxygen demand equivalents, $COD_{sol}$) in the influent and effluent. Black continuous line: average DO during the different representative periods, values on right axis.

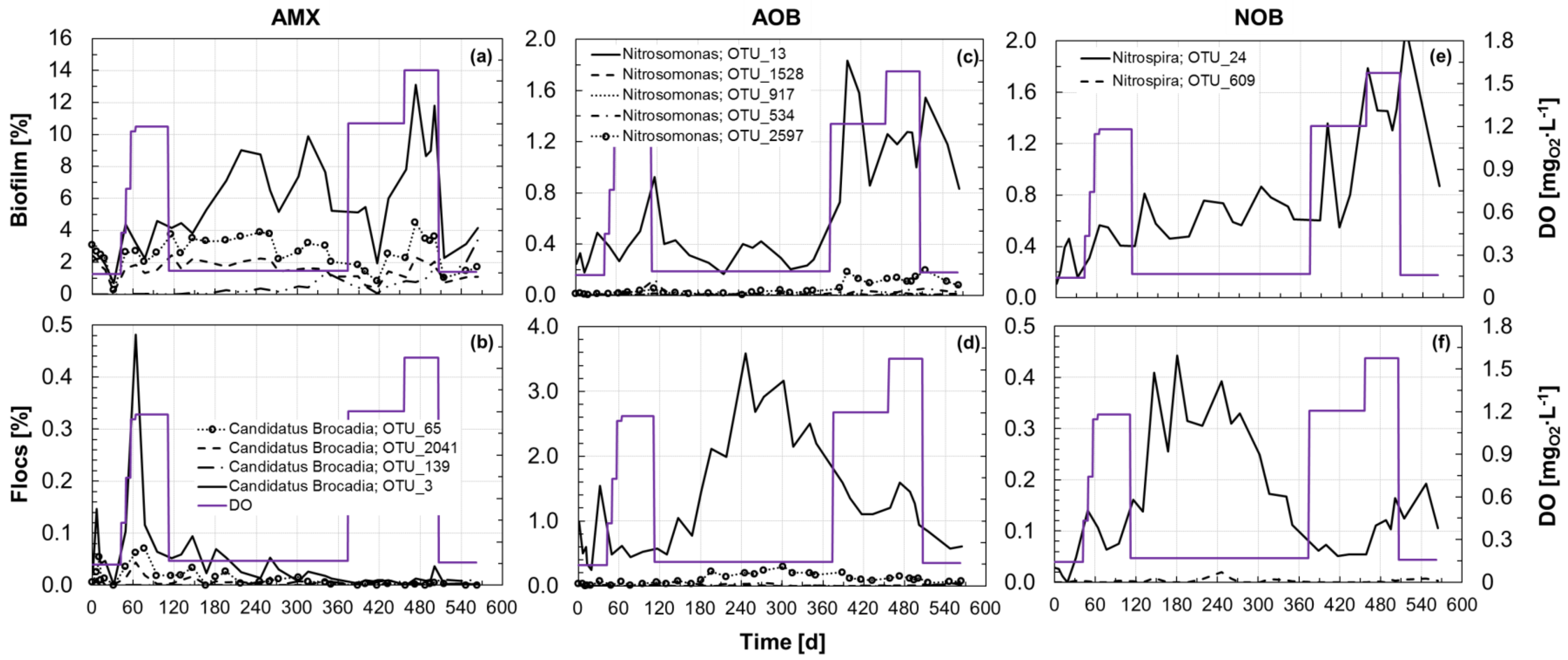


**Figure S4:** Time series of the relative abundance of the individual operational taxonomic units (OTUs) of the AMX (**a, b**), AOB (**c, d**) and NOB (**e, f**) guilds in the biofilm and flocs biomass fractions as estimated by 16S rRNA gene-based amplicon sequencing analysis. The average DO during the different representative periods is reported in all panels for reference in violet.

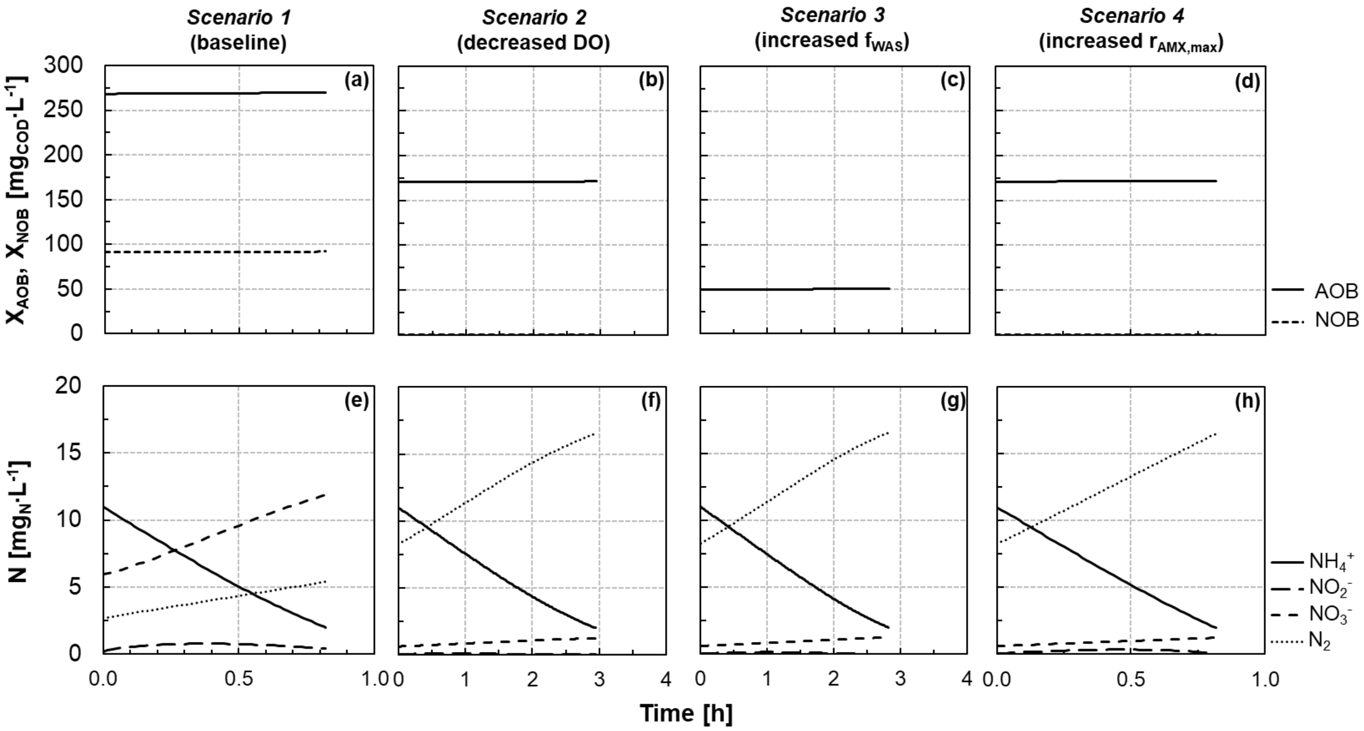


**Figure S5:** Simulated nitrogen species and biomass concentrations during an SBR cycle at pseudo steady-state for the four scenarios (see Figure 4).

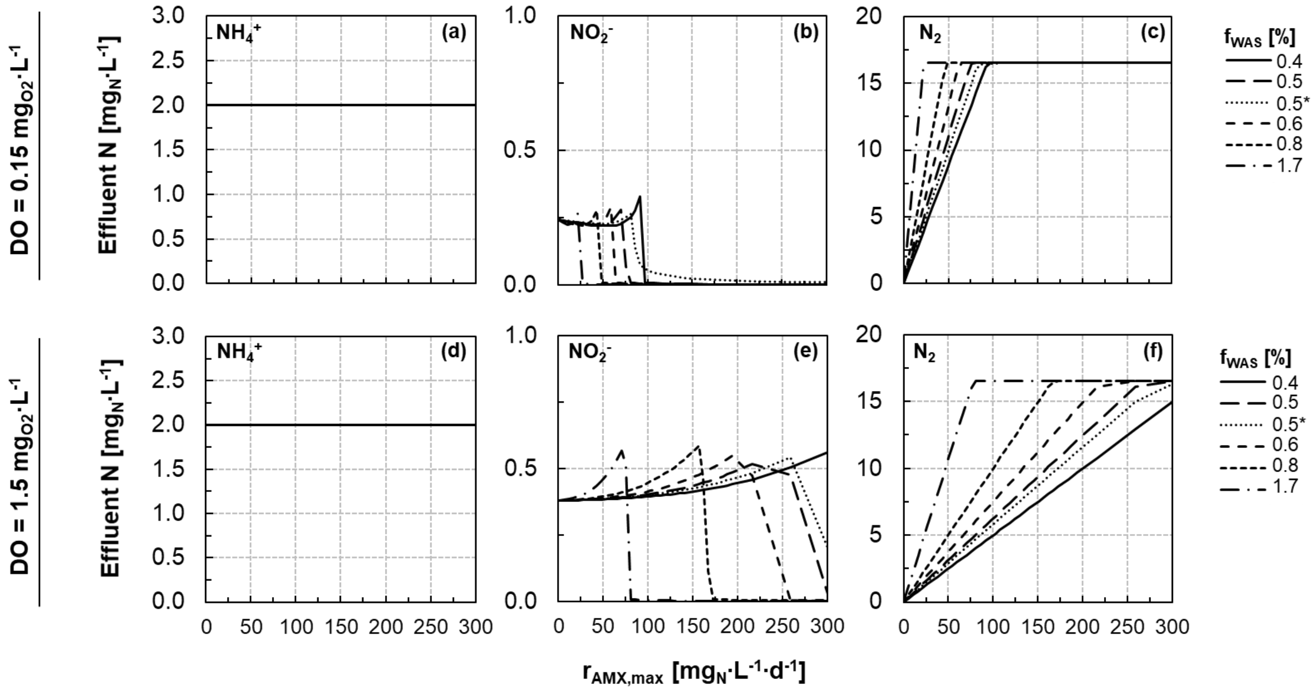


**Figure S6:** Simulated effluent concentrations of dissolved ammonium (**a, d**), nitrite (**b, e**) and di-nitrogen (**c, f**) at pseudo steady-state for the two reference DO concentrations, 0.15 and 1.5 $mg_{O2}·L^{-1}$ (see Figure 5). The different lines represent distinct fractions of flocs removed at the end of each cycle ($f_{WAS}$). $r_{AMX,max}$ is expressed as $mg_{(NH4+NO2)-N}·L^{-1}·d^{-1}$.

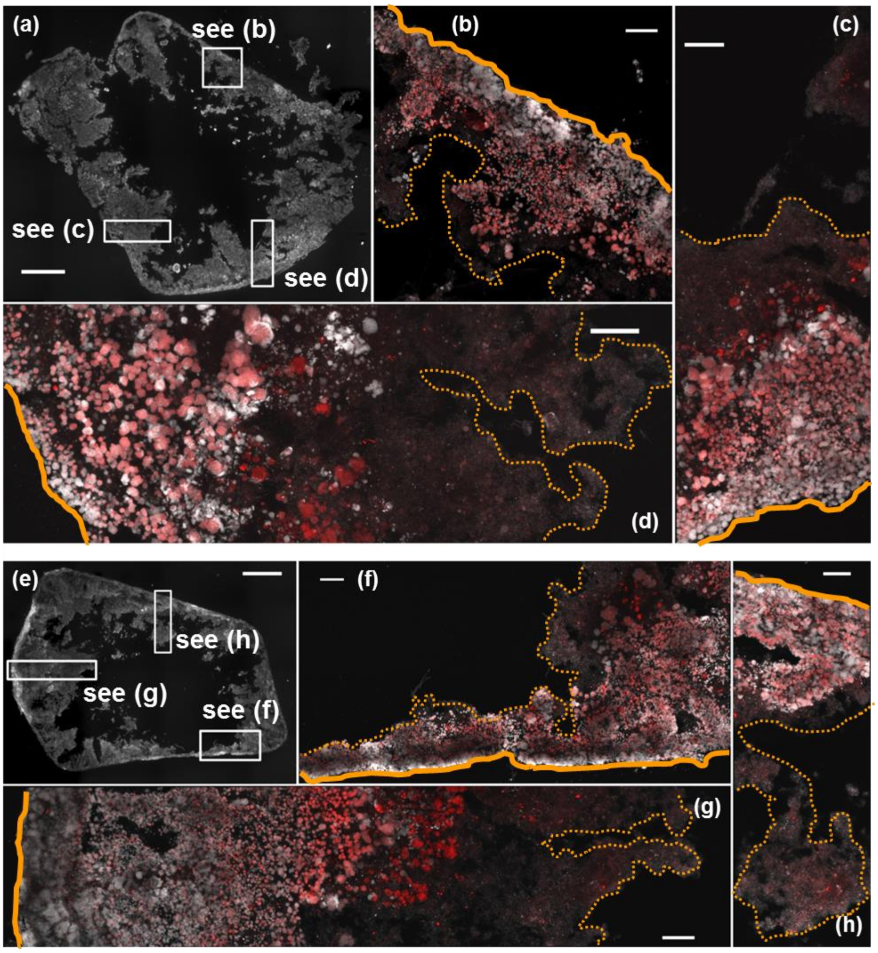


**Figure S7:** Thin-section FISH-stained micrographs of two biofilm carriers collected on day 130 (**a-d**) and 320 (**e-h**). (**a, e**) Wide-field epifluorescence micrograph showing DAPI-stained bacteria (white). Scale bar = 500 µm. (**b-d**, **f-h**) Confocal maximum intensity projections showing anammox bacteria (red) and DAPI-stained bacteria (white). Scale bars = 50 µm. *Continuous yellow line*: surface of the plastic carrier. *Dashed yellow line*: biofilm surface.

Method: Confocal microscopy was performed on K5 carrier cryosections using fluorescent *in situ* hybridization (FISH) probe mixes targeting anammox bacteria (Cy5 fluorophore; Amx 820 (Schmid *et al.*, 2000), and Bfu 613 (Kartal *et al.*, 2008)). K5 carrier biofilms were chemically fixed in a 4% formaldehyde solution for four hours, embedded in optical cutting temperature compound, and cryosectioned at -25°C to a thickness of 20 µm. Cryosections were hybridized overnight at 46°C in a 35% formamide v/v hybridization buffer containing probe concentrations of 0.5 µM. Hybridized cryosections were counterstained with 1 µg DAPI/mL in a glycerol-based antifadent mounting media. Image stacks were acquired on an inverted confocal microscope (Model TCS SP5, Leica Microsystems) equipped with an oil immersion 63x (1.44 NA) objective at a lateral resolution of 0.48 µm and an axial step size of 1 µm. DAPI and Cy5 fluorophores were excited sequentially with 405 nm and 633 nm laser lines, respectively.

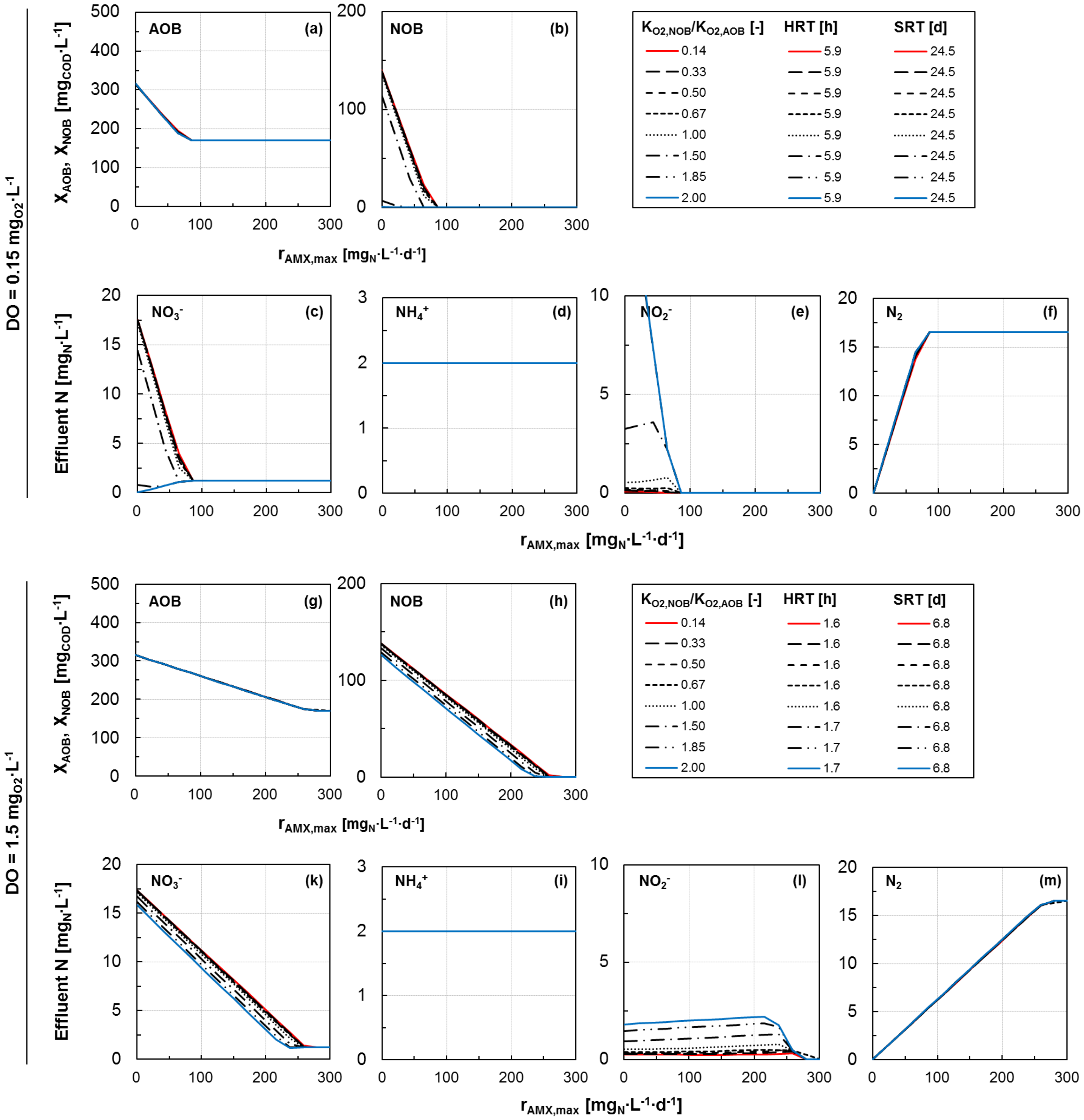


**Figure S8:** Sensitivity analysis of the model outputs to changes in $K_{O2,NOB}$/ $K_{O2,AOB}$. The considered values of the oxygen affinity constants for NOB and AOB are presented in Table S1, while the corresponding value of their ratio and the resulting HRT and SRT values at pseudo steady-state are reported in the legend. A flocs removal of 0.5% per cycle was assumed. DO of 0.15 $mg_{O2}\cdot L^{-1}$ (panels **a-f**) and 1.5 $mg_{O2}\cdot L^{-1}$ (panels **g-m**). (**a, g**) modelled pseudo steady-state AOB biomass concentration; (**b, h**) modelled pseudo steady-state NOB biomass concentration; (**c-f** and **k-m**) effluent $NO_3^-$, $NH_4^+$, $NO_2^-$ and $N_2$ concentrations. Note the different scales of the vertical axes in the graphs. $r_{AMX,max}$ is expressed as $mg_{(NH4+NO2)\text{-}N}\cdot L^{-1}\cdot d^{-1}$.

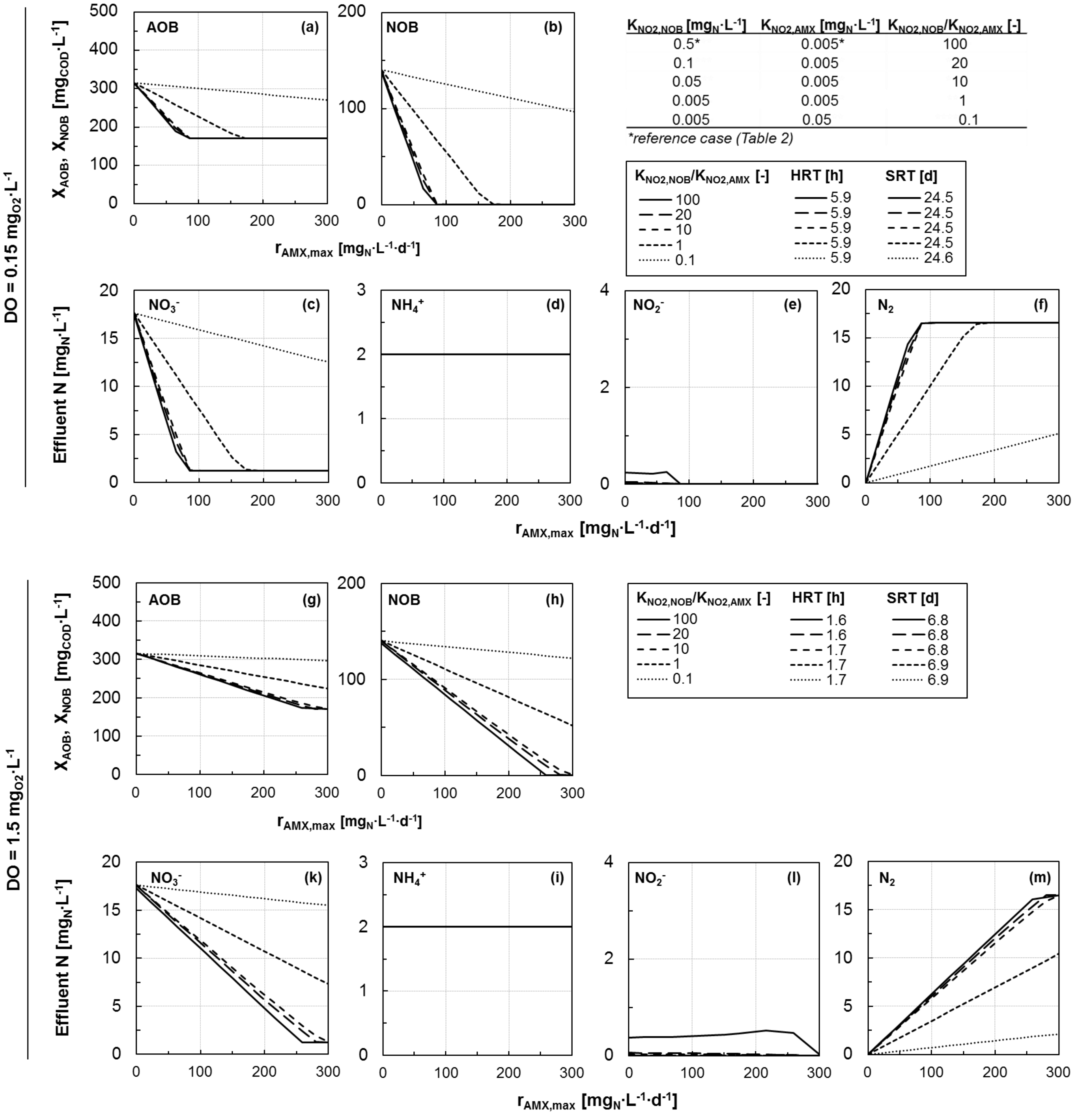


**Figure S9:** Sensitivity analysis of the model outputs to changes in $K_{NO2,NOB}$/ $K_{NO2,AMX}$. The considered values of the nitrite affinity constant for NOB and AMX are presented in the inserted Table S1. The corresponding value of their ratio and the resulting HRT and SRT values at pseudo steady-state are presented in the legend. *Reference scenario as in Table 2. A flocs removal of 0.5% per cycle was assumed. DO of 0.15 $mg_{O2}\cdot L^{-1}$ (panels **a-f**) and 1.5 $mg_{O2}\cdot L^{-1}$ (panels **g-m**). (**a, g**) modelled pseudo steady-state AOB biomass concentration; (**b, h**) modelled pseudo steady-state NOB biomass concentration; (**c-f** and **k-m**) effluent $NO_3^-$, $NH_4^+$, $NO_2^-$ and $N_2$ concentrations. Note the different scales of the vertical axes in the graphs. $r_{AMX,max}$ is expressed as $mg_{(NH4+NO2)-N}\cdot L^{-1}\cdot d^{-1}$.
