## Supplementary material for "Biomass segregation between biofilm and flocs improves the control of nitrite-oxidizing bacteria in mainstream partial nitritation and anammox processes"

### README.txt

```
PNAsim - Partial Nitrification/Anammox simulation
by Michele Laureni and Kris Villez

Website:       http://homepages.eawag.ch/~villezkr/spike/
Contact:      

This toolbox is a collection of Matlab functions and scripts originally written to simulate a simple model of a
hybrid MBBR operated for Partial Nitrification/Anammox process and with perfect biomass segregation (AOB and NOB
in the flocs and AMX in the biofilm). Sequencing batch mode operation is simulated, with a range of fixed
fractions of flocs removed per cycle and fixed active anammox biomass concentrations.

This collection is released to the public under the GPL v3 license in order to encourage the sharing of code and
prevent duplication of effort. If you find these functions useful, drop us a line. We also appreciate bug
reports and suggestions for improvements.

For more detail regarding copyrights see the file LICENSE.TXT located in the same folder where this README file
is located.

These files are provided as is, with no guarantees and are intended for non-commercial use.
These routines have been developed and tested successfully with Matlab (R2014b) on a Windows system.

To install the Matlab functions, follow the instructions below.

---------------------------------------------------
 I. Installation
---------------------------------------------------

1. Create a directory to contain all required toolboxes.  For example, in my case all toolbox directories are
in:

               C:\Tools\

2. Move the unzipped SCS folder (in which this README_SCS.txt file is located) to the following location:

               C:\Tools\PNAsim


-----------------------
 II. Using the toolbox
-----------------------

1. Start Matlab

2. To reproduce all results, execute the script

               RunScenarios_SBR.m

       in the folder

               C:\Tools\PNAsim\Application

----------------------------
 III. Relevant publications
----------------------------

Laureni et al., 2018 (submitted), "Biomass segregation between biofilm and flocs improves the control of nitrite
oxidizing bacteria in mainstream partial nitritation and anammox"

-------------
 IV. Changes
-------------

None reported yet.

------------------
 Acknowledgements
------------------


----------------------------------------------------------
Last modified on 27 November 2018
```

### TANbelow2

```
function [value,isterminal,direction] = TANbelow2(t,c)

% -------------------------------------------------------------------------
% Partial Nitrification/Anammox simulation - TANbelow2.m
% -------------------------------------------------------------------------
% Description
%
% TANbelow2 is a function which evaluates whether the minimal value of 2 is
% reached for the first element of the state vector.
%
% -------------------------------------------------------------------------
% Syntax
%
% I/O:  [value,isterminal,direction] = TANbelow2(t,c)
%
% INPUT
%       t           Time
%       c           State vector
%
% OUTPUT
%   value       Difference between c(1) and critical value of 2.
%   isterminal  Boolean, fixed to true
%   direction   Direction of check, fixed to negative (positive evaluation
%               if x(1) is below 2).
%
% -------------------------------------------------------------------------
% Last modification: Michele Laureni, 2018-11-27
% -------------------------------------------------------------------------

% -------------------------------------------------------------------------
% Copyright 2016 Michele Laureni and Kris Villez
%
% This file is part of a study in Matlab/Octave:
% Laureni et al., 2017 (submitted),
% " Biomass segregation between biofilm and flocs improves the
% control of nitrite-oxidizing bacteria in mainstream partial
% nitritation and anammox processes"
%
% This program is free software: you can redistribute it and/or modify it
% under the terms of the GNU General Public License as published by the
% Free Software Foundation, either version 3 of the License, or (at your
% option) any later version.
%
% This program is distributed in the hope that it will be useful, but
% WITHOUT ANY WARRANTY; without even the implied warranty of
% MERCHANTABILITY or FITNESS FOR A PARTICULAR PURPOSE.  See the GNU General
% Public License for more details.
%
% You should have received a copy of the GNU General Public License along
% with this program. If not, see <http://www.gnu.org/licenses/>.
% -------------------------------------------------------------------------


% Locate the time when height passes through zero in a decreasing direction
% and stop integration.
Value           =       c(1)-2  ;      % detect y-1/2 = 0
isterminal      =       1       ;      % stop the integration
direction       =       -1      ;      % negative direction
```

#### DefineSimulation.m

```
% ------------------------------------------------------------------------
% Partial Nitrification/Anammox simulation - DefineSimulation.m
% ------------------------------------------------------------------------
% Description
%
% DefineSimulation is a script which defines all the parameter
% for the simulation of a PN/A process operated in sequencing batch mode.
%
% ------------------------------------------------------------------------
% Last modification: Michele Laureni, 2018-11-27
% ------------------------------------------------------------------------

% ------------------------------------------------------------------------
% Copyright 2016 Michele Laureni and Kris Villez
%
% This file is part of a study in Matlab/Octave:
% Laureni et al., 2017 (submitted),
% "Biomass segregation between biofilm and flocs improves the control
% of nitrite oxidizing bacteria in mainstream partial nitritation
% and anammox"
%
% This program is free software: you can redistribute it and/or modify it
% under the terms of the GNU General Public License as published by the
% Free Software Foundation, either version 3 of the License, or (at your
% option) any later version.
%
% This program is distributed in the hope that it will be useful, but
% WITHOUT ANY WARRANTY; without even the implied warranty of
% MERCHANTABILITY or FITNESS FOR A PARTICULAR PURPOSE.  See the GNU General
% Public License for more details.
%
% You should have received a copy of the GNU General Public License along
% with this program. If not, see <http://www.gnu.org/licenses/>.
% ------------------------------------------------------------------------

% Fixed simulation parameters

TcycleMAX      =      100    ;      % d - maximal length for dynamically controlled cycle
Tcycle         =      6/24   ;      % d - length for fixed cycle
nCycleMax      =      1e6    ;

% SBRcontrol   =      'fixedrecipe'         ;
SBRcontrol     =      'dynamicrecipe'       ;

% 1. concentrations
cTAN_in        =      20     ;   % gN/m3
cTNO2_in       =      0      ;   % gN/m3
cTNO3_in       =      0      ;   % gN/m3
cN2_in         =      0      ;   % gN/m3
cAOB_in        =      300    ;   % gCOD/m3
cNOB_in        =      300    ;   % gCOD/m3

cAOB_0         =      300    ;   % gCOD/m3
cNOB_0         =      300    ;   % gCOD/m3
S_O2           =      0.15   ;   % gO2/m3


% Parameters in 2. and 3. from Laureni et al., 2017 (submitted):
% "Biomass segregation between biofilm and flocs improves the control
% of nitrite oxidizing bacteria in mainstream partial nitritation
% and anammox"

% 2. stoichiometrix matrix
iN_AOB         =      0.083  ;   % gN/gCOD
iN_NOB         =      0.083  ;   % gN/gCOD
iN_AMX         =      0.058  ;   % gN/gCOD
Y_AOB          =      0.18   ;   % gCOD/gN
Y_NOB          =      0.08   ;   % gCOD/gN
Y_AMX          =      0.17   ;   % gCOD/gN

% 3. reaction rates
mu_AOB         =      0.297  ;   % 1/d
mu_NOB         =      0.337  ;   % 1/d
```

```
mu_AMX          =       0.017   ;   % 1/d

b_AOB           =       0       ;       % 1/d
b_NOB           =       0       ;       % 1/d

K_AOB_NH4       =       2.4     ;   % gN/m3
K_AMX_NH4       =       0.03    ;   % gN/m3

K_NOB_NO2       =       0.5     ;   % gN/m3
K_AMX_NO2       =       0.005   ;   % gN/m3

K_AOB_O2        =       0.6     ;   % gCOD/m3
K_NOB_O2        =       0.4     ;   % gCOD/m3

% Model structure
N       =   [  -1/Y_AOB-iN_AOB       +1/Y_AOB            0           0           1              0        ;
               -iN_NOB               -1/Y_NOB            +1/Y_NOB    0           0              1        ;
               -1/Y_AMX-iN_AMX       -1/Y_AMX-1/1.14     +1/1.14     2/Y_AMX     0              0        ;
               +iN_AOB               0                   0           0           -1             0        ;
               +iN_NOB               0                   0           0           0              -1      ];
            %  gN/gCOD               gN/gCOD             gN/gCOD     gN/gCOD     gCOD/gCOD      gCOD/gCOD

iTAN            =       1       ;
iTNO2           =       2       ;
iTNO3           =       3       ;
iN2             =       4       ;
iAOB            =       5       ;
iNOB            =       6       ;
nC              =       6       ; % number of simulated concentrations

rho     =   @(t,c,X_AMX,S_O2) [     ...
    mu_AOB     .*c(iAOB,:)    .*c(iTAN,:)./(c(iTAN,:)+K_AOB_NH4)          .*S_O2/(S_O2+K_AOB_O2)               ;
%   1/d        gCOD/m3        (gN/m3) / ( gN/m3 + gN/m3 )                 (gCOD/m3) / ( gCOD/m3 + gCOD/m3 )
    mu_NOB     .*c(iNOB,:)    .*c(iTNO2,:)./(c(iTNO2,:)+K_NOB_NO2)        .*S_O2/(S_O2+K_NOB_O2)               ;
%   1/d        gCOD/m3        (gN/m3) / ( gN/m3 + gN/m3 )                 (gCOD/m3) / ( gCOD/m3 + gCOD/m3 )
    mu_AMX     .*X_AMX        .*c(iTAN,:)./(c(iTAN,:)+K_AMX_NH4)          .*c(iTNO2,:)./(c(iTNO2,:)+K_AMX_NO2) ;
%   1/d        gCOD/m3        (gN/m3) / ( gN/m3 + gN/m3 )                 (gN/m3) / ( gN/m3 + gN/m3 )
    b_AOB      .*c(iAOB,:)                                                                                     ;
%   1/d        gCOD/m3
    b_NOB      .*c(iNOB,:)                                                                  ]                  ;
%   1/d        gCOD/m3

% unit rho:    gCOD/m3.d
```

#### RunOneScenario_SBR.m

```
% --------------------------------------------------------------------------
% Partial Nitrification/Anammox simulation - RuneOneScenario_SBR.m
% --------------------------------------------------------------------------
% Description
%
% RuneOneScenario_SBR is a script which simulates a hybrid MBBR operated for
% PN/A in sequencing batch mode. Perfect biomass segregation with AOB and
% NOB in the flocs and AMX in the biofilm, is assumed. One fixed
% fraction of flocs removed per cycle and one fixed active anammox biomass
% concentration are simulated.
%
% --------------------------------------------------------------------------
% Last modification: Michele Laureni, 2018-11-27
% --------------------------------------------------------------------------

% --------------------------------------------------------------------------
% Copyright 2016 Michele Laureni and Kris Villez
%
% This file is part of a study in Matlab/Octave:
% Laureni et al., 2017 (submitted),
% "Biomass segregation between biofilm and flocs improves the control
% of nitrite oxidizing bacteria in mainstream partial nitritation
% and anammox"
%
% This program is free software: you can redistribute it and/or modify it
% under the terms of the GNU General Public License as published by the
% Free Software Foundation, either version 3 of the License, or (at your
% option) any later version.
%
% This program is distributed in the hope that it will be useful, but
% WITHOUT ANY WARRANTY; without even the implied warranty of
% MERCHANTABILITY or FITNESS FOR A PARTICULAR PURPOSE.  See the GNU General
% Public License for more details.
%
% You should have received a copy of the GNU General Public License along
% with this program. If not, see <http://www.gnu.org/licenses/>.
% --------------------------------------------------------------------------

% Initialize state vector
C               =       nan(nC,1)    ;
C(iTAN)         =       cTAN_in      ;
C(iTNO2)        =       cTNO2_in     ;
C(iTNO3)        =       cTNO3_in     ;
C(iN2)          =       cN2_in       ;
C(iAOB)         =       cAOB_0       ;
C(iNOB)         =       cNOB_0       ;

TimeNow         =       0            ;

iCycle          =       0            ;
TT              =       0            ;
CC              =       C(:)'        ;

BiomassRet      =       1-WAS        ;
deltaT          =       TcycleMAX    ;

while TT(end)<50*(-deltaT/log(BiomassRet))  &&  iCycle<nCycleMax

    if mod(iCycle,100)==0
        disp([ 'Simulation progress: ' num2str((TT(end)/50)/(-deltaT/log(BiomassRet))*100,'%3.1f') ' %' ]);
    end

    iCycle     =      iCycle+1                      ;

    % ======================================================================
    % Fill stage: reset initial species concentrations

    C(iTAN)    =      (C(iTAN) +  cTAN_in    )/2     ;
    C(iTNO2)   =      (C(iTNO2)+  cTNO2_in   )/2     ;
    C(iTNO3)   =      (C(iTNO3)+  cTNO3_in   )/2     ;
    C(iN2)     =      (C(iN2)  +  cN2_in     )/2     ;
    C(iAOB)    =      C(iAOB)                        ;
    C(iNOB)    =      C(iNOB)                        ;
```

```
    % ======================================================================
    % React stage: biochemical conversion

    switch SBRcontrol
        case 'fixedrecipe'
            [tsim,csim]=ode15s(                 ...
                @(t,c) N'*rho(t,c,X_AMX,S_O2),  ...     % Unit N'*rho: [ gN/m3.d gN/m3.d gCOD/m3.d gCOD/m3.d ]
                [0:Tcycle/100:Tcycle],          ...
                C );
        case 'dynamicrecipe'
            opts=odeset('Events',@TANbelow2);
            [tsim,csim]=ode15s( @(t,c)          ...
                N'*rho(t,c,X_AMX,S_O2),         ...     % Unit N'*rho: [ gN/m3.d gN/m3.d gCOD/m3.d gCOD/m3.d ]
                [0:(1/24/60):TcycleMAX],        ...
                C,                              ...
                opts);
    end

    C(:)        =     csim(end,:)'                ;

    % ======================================================================
    % Sedimentation/decant stage: adjust biomass concentrations

    deltaT      =     tsim(end)                   ;
    TimeNow     =     TimeNow+deltaT              ;

    C(iAOB)     =     C(iAOB)*BiomassRet          ;
    C(iNOB)     =     C(iNOB)*BiomassRet          ;

    TT          =     [   TT  ;      TimeNow     ]  ;
    CC          =     [   CC  ;      C(:)'       ]  ;


end

disp([ 'Simulation progress: ' num2str(100,'%3.1f') ' %' ]);
```

#### RunScenarios_SBR.m

```
% -------------------------------------------------------------------------
% Partial Nitrification/Anammox simulation - RunScenarios_SBR.m
% -------------------------------------------------------------------------
% Description
%
% RunScenarios_SBR is a script which simulates a hybrid MBBR operated for
% PN/A in sequencing batch mode. Perfect biomass segregation with AOB and
% NOB in the flocs and AMX in the biofilm, is assumed. A range of fixed
% fractions of flocs removed per cycle and of fixed active anammox biomass
% concentrations are simulated.
%
% -------------------------------------------------------------------------
% Last modification: Michele Laureni, 2018-11-27
% -------------------------------------------------------------------------

% -------------------------------------------------------------------------
% Copyright 2016 Michele Laureni and Kris Villez
%
% This file is part of a study in Matlab/Octave:
% Laureni et al., 2017 (submitted),
% "Biomass segregation between biofilm and flocs improves the control
% of nitrite oxidizing bacteria in mainstream partial nitritation
% and anammox"
%
% This program is free software: you can redistribute it and/or modify it
% under the terms of the GNU General Public License as published by the
% Free Software Foundation, either version 3 of the License, or (at your
% option) any later version.
%
% This program is distributed in the hope that it will be useful, but
% WITHOUT ANY WARRANTY; without even the implied warranty of
% MERCHANTABILITY or FITNESS FOR A PARTICULAR PURPOSE.  See the GNU General
% Public License for more details.
%
% You should have received a copy of the GNU General Public License along
% with this program. If not, see <http://www.gnu.org/licenses/>.
% -------------------------------------------------------------------------

clc
clear all
close all

% Scenario simulation parameters

vec_WAS         =       [ 4, 5, 6, 8, 17 ]/1000       ;        %   flocs fraction removed per cycle
vec_X_AMX       =       [ 0:100:1400 ]                ;        %   gCOD/m^3

% -------------------------------------------
% Initialization and pre-allocation

addpath('../Functions/')

DefineSimulation

nWAS            =       length(vec_WAS)       ;
nAMX            =       length(vec_X_AMX)     ;
CC_steady       =       nan(nWAS,nAMX,nC)     ;
SRT             =       nan(nWAS,nAMX)        ;

% -------------------------------------------
% Execute every scenario

for iAMX=1:nAMX
    for iWAS = 1:nWAS

        clc

        disp('Simulation')
        disp(['  AMX: ' num2str(iAMX) ' of ' num2str(nAMX) ])
        disp(['  Ret: ' num2str(iWAS) ' of ' num2str(nWAS) ])

        % Set simulation-specific parameters:
        WAS                     =    vec_WAS(iWAS)            ;
```

```
        X_AMX                   =    vec_X_AMX(iAMX)         ;

        % Execute simulation
        RunOneScenario_SBR

        % Get final concentration vector and store it one big array
        Cfinal                       =      CC(end,:)                   ;
        CC_steady(iWAS,iAMX,:)       =      Cfinal                      ;
        SRT(iWAS,iAMX)               =      -deltaT/log(BiomassRet)     ;

        save(['..\Results\Simulation_' num2str(iWAS) '_' num2str(iAMX)],'TT','CC','tsim','csim')

    end
end

close all
drawnow

% Make some plots

figure,
subplot(2,1,1), hold on
    plot(tsim,csim(:,iTAN),'.')
    plot(tsim,csim(:,iTNO2),'.')
    plot(tsim,csim(:,iTNO3),'.')
    plot(tsim,csim(:,iN2),'.')
    xlabel(['Time [d]'])
    ylabel(['Concentration [gN/m^3]'])
    legend({'TAN','TNO_2N','TNO_3N','N_2'})

    % excel: [time, TAN, TNO2, TNO3, N2, X_AOB, X_NOB]
    filename                    =    '..\Results\9_Cycle_N_X_time.xlsx'             ;
    MMM                         =   ones(size(csim,1),(size(csim,2)+1))             ;
    MMM(:,1)                    =   tsim.*24                                        ;
    MMM(:,2:(size(csim,2)+1))   =   csim                                            ;
    xlswrite(filename, MMM)

subplot(2,1,2), hold on
    plot(tsim,csim(:,iAOB),'.')
    plot(tsim,csim(:,iNOB),'.')
    xlabel(['Time [d]'])
    ylabel(['Concentration [gN/m^3]'])
    legend({'X_{AOB}','X_{NOB}'})

figure,
subplot(2,1,1), hold on
    plot(TT,CC(:,iTAN),'.')
    plot(TT,CC(:,iTNO2),'.')
    plot(TT,CC(:,iTNO3),'.')
    plot(TT,CC(:,iN2),'.')
    xlabel(['Time [d]'])
    ylabel(['Concentration [gN/m^3]'])
    legend({'TAN','TNO_2N','TNO_3N','N_2'})
    %set(gca,'Ylim',[0 2.1])

    % excel: [time, TAN, TNO2, TNO3, N2, X_AOB, X_NOB]
    filename                    =   '..\Results\10_Equilibrium_N_X_time.xlsx'       ;
    MMM                         =   ones(size(CC,1),(size(CC,2)+1))                 ;
    MMM(:,1)                    =   TT                                              ;
    MMM(:,2:(size(CC,2)+1))     =   CC                                              ;
    xlswrite(filename, MMM)

subplot(2,1,2),  hold on
    plot(TT,CC(:,iAOB),'.')
    plot(TT,CC(:,iNOB),'.')
    xlabel(['Time [d]'])
    ylabel(['Concentration [gN/m^3]'])
    legend({'X_{AOB}','X_{NOB}'})
    %set(gca,'Ylim',[0 100])
    drawnow

outcomename = {'TAN','TNO2','X AOB','X NOB'};


figure(174),
subplot(2,2,1), hold on
```

```
    plot(vec_X_AMX,squeeze(CC_steady(:,:,1)),'+-')  %S_NH4_X_anammox
    ylabel('TAN [g/m^3]')
    set(gca,'Ylim',[0 20])

    filename                    =   '..\Results\3_NH4_X_AMX.xlsx'                ;
    S_NH4                       =   squeeze(CC_steady(:,:,1))                    ;
    MMM                         =   ones((size(vec_WAS,2)+1),(size(vec_X_AMX,2)+1)) ;
    MMM(1,:)                    =   transpose([0;transpose(vec_X_AMX)])          ;
    MMM(:,1)                    =   transpose([0;transpose(vec_WAS).*100])       ;
    MMM(2:(size(vec_WAS,2)+1),2:(size(vec_X_AMX,2)+1))  ...
                                =   S_NH4                                        ;
    xlswrite(filename, transpose(MMM))

subplot(2,2,2), hold on
    plot(vec_X_AMX,squeeze(CC_steady(:,:,2)),'+-')  %NO2_X_AMX
    ylabel('TNO2N [g/m^3]')
    set(gca,'Ylim',[0 20])
    filename                    =   '..\Results\4_NO2_X_AMX.xlsx'                ;
    S_NO2                       =   squeeze(CC_steady(:,:,2))                    ;
    MMM                         =   ones((size(vec_WAS,2)+1),(size(vec_X_AMX,2)+1)) ;
    MMM(1,:)                    =   transpose([0;transpose(vec_X_AMX)])          ;
    MMM(:,1)                    =   transpose([0;transpose(vec_WAS).*100])       ;
    MMM(2:(size(vec_WAS,2)+1),2:(size(vec_X_AMX,2)+1))  ...
                                =   S_NO2                                        ;
    xlswrite(filename, transpose(MMM))

subplot(2,2,3), hold on
    plot(vec_X_AMX,squeeze(CC_steady(:,:,3)),'+-')  %NO3_X_AMX
    ylabel('TNO3N [g/m^3]')
    xlabel('X_{AMX} [g/m^3]')
    set(gca,'Ylim',[0 20])

    filename                    =   '..\Results\5_NO3_X_AMX.xlsx'                ;
    S_NO3                       =   squeeze(CC_steady(:,:,3))                    ;
    MMM                         =   ones((size(vec_WAS,2)+1),(size(vec_X_AMX,2)+1)) ;
    MMM(1,:)                    =   transpose([0;transpose(vec_X_AMX)])          ;
    MMM(:,1)                    =   transpose([0;transpose(vec_WAS).*100])       ;
    MMM(2:(size(vec_WAS,2)+1),2:(size(vec_X_AMX,2)+1))  ...
                                =   S_NO3                                        ;
    xlswrite(filename, transpose(MMM))

subplot(2,2,4), hold on
    plot(vec_X_AMX,squeeze(CC_steady(:,:,4)),'+-')  %N2_X_AMX
    ylabel('6_N2 [g/m^3]')
    legend([cellstr(num2str(vec_WAS(:)*100)) ])
    set(gca,'Ylim',[0 20])

    filename                    =   '..\Results\6_N2_X_AMX.xlsx'                 ;
    S_N2                        =   squeeze(CC_steady(:,:,4))                    ;
    MMM                         =   ones((size(vec_WAS,2)+1),(size(vec_X_AMX,2)+1)) ;
    MMM(1,:)                    =   transpose([0;transpose(vec_X_AMX)])          ;
    MMM(:,1)                    =   transpose([0;transpose(vec_WAS).*100])       ;
    MMM(2:(size(vec_WAS,2)+1),2:(size(vec_X_AMX,2)+1))  ...
                                =   S_N2                                         ;
    xlswrite(filename, transpose(MMM))

figure(374),
subplot(2,2,1), hold on
    plot(vec_X_AMX,squeeze(CC_steady(:,:,5))','+-')
    ylabel('X_{AOB} [g/m^3]')
    set(gca,'Xtick',[0:100:1000])

    filename                    =   '..\Results\1_AOB_X_AMX.xlsx'                ;
    AOB                         =   squeeze(CC_steady(:,:,5))                    ;
    MMM                         =   ones((size(vec_WAS,2)+1),(size(vec_X_AMX,2)+1)) ;
    MMM(1,:)                    =   transpose([0;transpose(vec_X_AMX)])          ;
    MMM(:,1)                    =   transpose([0;transpose(vec_WAS).*100])       ;
    MMM(2:(size(vec_WAS,2)+1),2:(size(vec_X_AMX,2)+1))  ...
                                =   AOB                                          ;
    xlswrite(filename, transpose(MMM))

subplot(2,2,2), hold on
    plot(vec_X_AMX,squeeze(CC_steady(:,:,6))','+-')
    ylabel('X_{NOB} [g/m^3]')
    set(gca,'Xtick',[0:100:1000])
%     set(gca,'Ylim',[0 150])
```

```
    filename                        =   '..\Results\2_NOB_X_AMX.xlsx'                       ;
    NOB                             =   squeeze(CC_steady(:,:,6))                           ;
    MMM                             =   ones((size(vec_WAS,2)+1),(size(vec_X_AMX,2)+1))     ;
    MMM(1,:)                        =   transpose([0;transpose(vec_X_AMX)])                 ;
    MMM(:,1)                        =   transpose([0;transpose(vec_WAS).*100])              ;
    MMM(2:(size(vec_WAS,2)+1),2:(size(vec_X_AMX,2)+1))  ...
                                    =   NOB                                                 ;
    xlswrite(filename, transpose(MMM))

subplot(2,2,3), hold on
    plot(vec_X_AMX,(-SRT.*repmat(log(1-vec_WAS(:)),[1 nAMX]))','+-')
    ylabel('Cycle length [d]')
    xlabel('X_{AMX} [g/m^3]')
    set(gca,'Xtick',[0:100:1000])

    filename                        =   '..\Results\7_HRT_X_AMX.xlsx'                       ;
    Cycle_Length                    =   -SRT.*repmat(log(1-vec_WAS(:)),[1 nAMX])            ;
    MMM                             =   ones((size(vec_WAS,2)+1),(size(vec_X_AMX,2)+1))     ;
    MMM(1,:)                        =   transpose([0;transpose(vec_X_AMX)])                 ;
    MMM(:,1)                        =   transpose([0;transpose(vec_WAS).*100])              ;
    MMM(2:(size(vec_WAS,2)+1),2:(size(vec_X_AMX,2)+1))  ...
                                    =   Cycle_Length.*(2*24)                                ;
    xlswrite(filename, transpose(MMM))

subplot(2,2,4), hold on
    plot(vec_X_AMX,(+SRT)','+-')
    xlabel('X_{AMX} [g/m^3]')
    ylabel('SRT [d]')
    set(gca,'Xtick',[0:100:1000])
    legend([cellstr(num2str(vec_WAS(:)*100)) ])

    filename                        =   '..\Results\8_SRT_X_AMX.xlsx'                       ;
    SRT_final                       =   squeeze((+SRT))                                     ;
    MMM                             =   ones((size(vec_WAS,2)+1),(size(vec_X_AMX,2)+1))     ;
    MMM(1,:)                        =   transpose([0;transpose(vec_X_AMX)])                 ;
    MMM(:,1)                        =   transpose([0;transpose(vec_WAS).*100])              ;
    MMM(2:(size(vec_WAS,2)+1),2:(size(vec_X_AMX,2)+1))  ...
                                    =   SRT_final                                           ;
    xlswrite(filename, transpose(MMM))
```
